# Paired RNA profiling of circulating small extracellular vesicles links survival in ALS to reactive glial - vascular programs

**DOI:** 10.64898/2026.09.08.749915

**Authors:** Jonathan S. Weerakkody, Finja Bokstaller, Uma Sthanu, David Pitt

## Abstract

Circulating RNA can capture molecular variation associated with amyotrophic lateral sclerosis (ALS) progression, but biofluid heterogeneity makes biologically organized signals difficult to recover. Here, we paired microRNA (miRNA) and messenger RNA (mRNA) profiles from glutamate–aspartate transporter (GLAST)-positive small extracellular-vesicles (sEVs) to estimate survival time and identify survival-associated programs. In 45 participants with ALS and 15 controls, biologically constrained multi-omic factor analysis identified a program in which reciprocal miRNA–mRNA states among target-supported pairs tracked survival. The miRNA arm was then evaluated in an independent total-plasma cohort of 248 participants with ALS, refining a five-miRNA panel that added prognostic information beyond functional decline and neurofilament light chain, particularly over longer survival horizons. The mRNA arm was evaluated across 586 cortical profiles from 308 donors and 527,261 nuclei from 69 donors, identifying a five-gene core associated with glial reactivity, vascular programs and reduced myelinating identity. Together, these findings establish a framework for integrating regulatory RNA layers in circulation to identify clinically relevant programs and relate them to disease-relevant cellular states.

## Introduction

Amyotrophic lateral sclerosis (ALS) is marked by substantial clinical and biological heterogeneity, with pronounced variation in the rate of functional decline and survival.^1–4^ This variability complicates individual prognosis and increases outcome variance in clinical trials, reducing their precision and interpretability.^5–7^ Biomarkers that capture underlying disease biology could therefore improve risk stratification and trial design. Neurofilament light chain (NfL), a marker of neuroaxonal injury, is the most established blood biomarker in ALS and provides robust prognostic, pharmacodynamic and disease-stage information.^7–14^ Recent longitudinal studies further indicate that circulating biomarker trajectories are temporally heterogeneous across the ALS disease course, motivating complementary molecular readouts of disease state.^9,15,16^ Glial, immune and vascular changes may precede or accompany neuroaxonal injury, and disease progression reflects coordinated dysfunction across astrocytic,^17–19^ microglial,^20^ oligodendroglial^21^ and neurovascular compartments.^22^ Human post-mortem transcriptomics resolves this multicellular pathology into cortical states associated with molecular subtype and prognosis,^23–25^ but relating circulating biomarkers to these tissue-resolved programs remains difficult.

Circulating RNA offers a potential source of biologically informative signals that complement measures of tissue injury. ^26–31^ The challenge is that RNA in biofluids is intrinsically heterogeneous: transcripts originate from multiple tissues and are distributed among extracellular vesicles, protein-bound complexes, and lipoprotein-associated particles.^32–35^ Disease-relevant central nervous system (CNS) RNA is therefore embedded within a much larger molecular background, creating both biological and statistical noise. This is evident in high-dimensional biofluid datasets where dominant tissue sources, technical effects, and cohort-specific structure can account for more variance than the progression-linked signal itself, causing unconstrained discovery to favor statistically prominent features that are not necessarily the most biologically informative. This may contribute to the limited convergence of candidate biomarkers across systematic liquid-biopsy^36^ and extracellular-vesicle RNA studies.^37^ The central challenge is therefore to move past identifying circulating associations and to recover biologically organized signals within them, then assign those signals interpretable context.

We reasoned that biologically constraining the search space before statistical ranking could recover more disease-relevant signals. The broader premise is that paired miRNA and mRNA measurements can reveal clinically relevant molecular programs even when they explain little overall variance; here, we tested this approach in ALS. Surface-selected extracellular vesicles (EVs) enrich a defined circulating RNA compartment while retaining selectively packaged cargo. Within this cargo, miRNAs and mRNAs provide an additional layer of biological organization through established post-transcriptional relationships: miRNAs represent a regulatory layer, whereas mRNAs report transcript abundance in the same preparation. We implemented this framework using GLAST (glutamate–aspartate transporter; excitatory amino-acid transporter 1, encoded by SLC1A3)-positive small extracellular vesicles (sEVs), an astrocyte-enriched circulating compartment previously identified in ALS.^38,39^ Established extracellular-vesicle (EV) RNA sorting mechanisms^40–43^ and experimentally supported miRNA–mRNA relationships^44–47^ provided independent biological priors for interpreting how the recovered cargo is packaged and directionally organized.

Here, we tested this paired-RNA strategy through a primary discovery analysis followed by two orthogonal external analyses. We first asked whether biologically constrained paired-RNA profiling could identify a survival-associated program organized by directional miRNA–mRNA relationships and not by statistical prominence alone, testing this in GLAST-positive sEVs from 45 participants with ALS and 15 controls. We then asked whether the miRNA arm remained measurable and prognostically informative in the more heterogeneous setting of unfractionated total plasma, using an independent cohort of 248 participants with ALS.^31^ In parallel, we evaluated the nominated mRNA arm across 586 human cortical transcriptomes^24^ and 527,261 nuclei.^25^ to determine whether these circulating candidates were represented within disease-relevant tissue programs. The plasma analysis therefore tested cross-compartment transfer of the miRNA signal, whereas the cortical analyses provided independent biological context for the mRNA candidates. Together, this design identifies a survival-associated paired-RNA program in circulation and tests its transferability and biological context in independent resources.

## Results

### GLAST-positive sEVs support paired profiling of regulatory and effector RNA

GLAST immunocapture recovered a rare, surface-defined sEV fraction from 0.5 ml plasma (Fig. 1a; Supplementary Note 1). Nanoparticle tracking of ten matched characterization preparations yielded a mean particle diameter of 138.8 ± 8.9 nm and a mean captured-to-total particle concentration ratio of 0.180% (median 0.124%; Fig. 1b). Immunogold electron microscopy localized GLAST to vesicle-sized structures (Fig. 1c), while three-colour direct stochastic optical reconstruction microscopy (dSTORM) resolved heterogeneous CD9, CD63 and CD81 phenotypes, including 12 triple-positive particles among 339 classified particles (3.5%; Fig. 1d and Extended Data Fig. 1a,b). Nano-flow cytometry provided complementary tetraspanin and size measurements (Extended Data Fig. 1a,d). Following Minimum Information for Studies of Extracellular Vesicles 2023 (MISEV2023) recommendations,^48^ we use sEV as an operational term for this size– and surface-defined fraction. Archived protein-assay comparisons showed predominantly positive GLAST-positive-to-total-EV ratios for GFAP, AQP4, GLT1/SLC1A2 and ALDOC (Fig. 1e). ApoB100 was lower in the GLAST-positive than GLAST-negative fraction in all three displayed records, and platelet CD41–CD61 was lower in two of three records (Fig. 1f). These preparations were used for paired-RNA discovery.

**Fig. 1.**
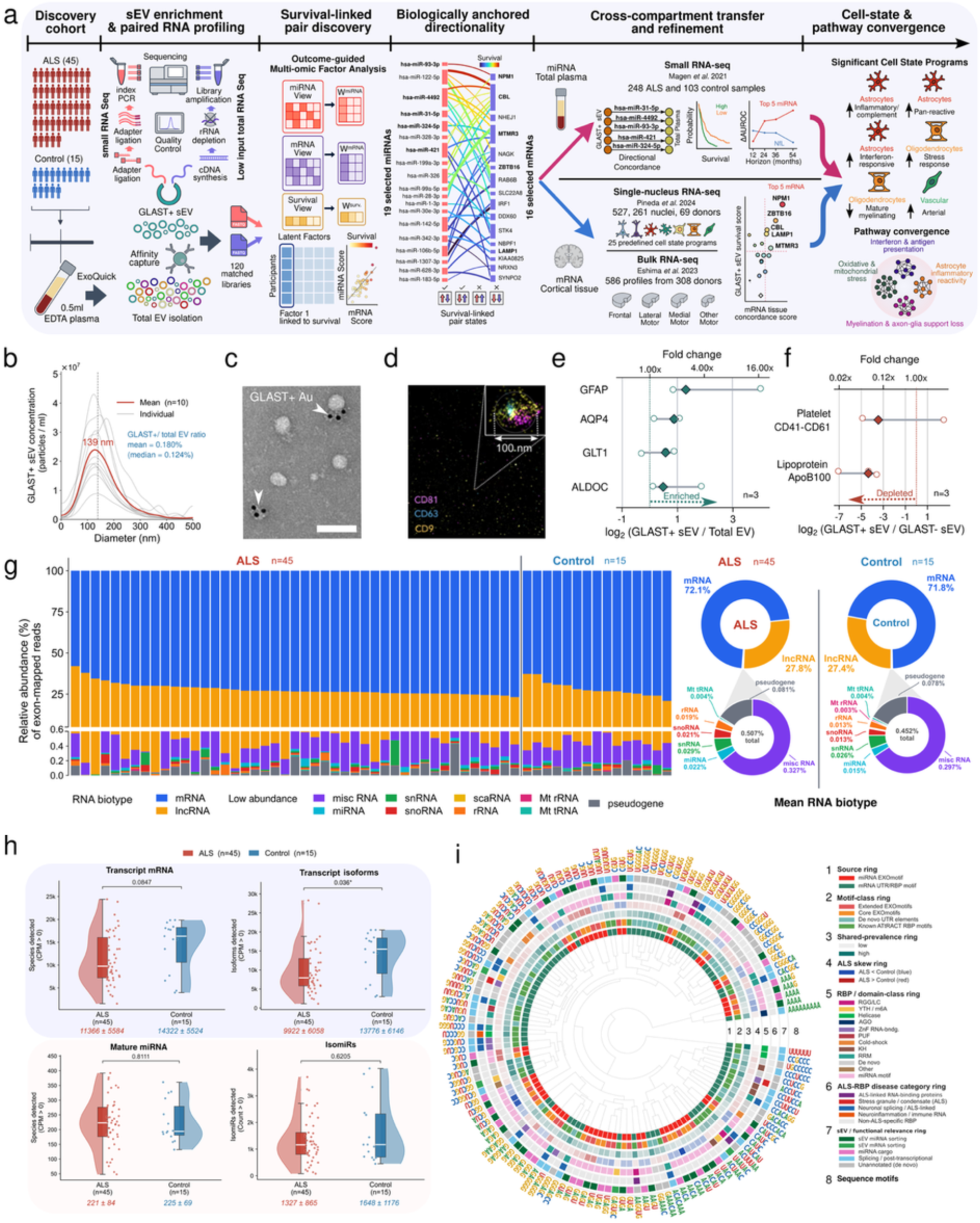
| GLAST enrichment defines a rare sEV compartment that preserves paired regulatory and effector RNA. **a**, Study design. GLAST/SLC1A3-positive sEVs were affinity-enriched from 0.5 ml plasma in 45 participants with ALS and 15 controls; one enriched RNA eluate from each participant generated a small-RNA and a low-input transcriptome library (120 matched libraries across 60 participants). The nominated candidates were subsequently evaluated and refined in Magen et al.^31^ total plasma (248 ALS, 103 controls), Eshima et al.^24^ bulk cortex (586 profiles from 308 donors) and Pineda et al.^25^ single-nucleus cortex (527,261 analyzed nuclei from 69 donors). **b,** Nanoparticle tracking of ten matched technical preparations, separate from sequenced participants. Grey curves, individual preparations; red, smoothed mean; dashed line, mean particle diameter (138.8 ± 8.9 nm). The captured-to-total particle concentration ratio averaged 0.180% (median 0.124%). **c,** Representative immunogold transmission electron micrograph; arrowheads indicate GLAST-labelled vesicle-sized structures. Scale bar, 200 nm. **d,** Three-colour dSTORM reconstruction of an individual particle showing CD9, CD63 and CD81; 12 of 339 dSTORM-classified particles were triple-positive (3.5%; rounded to 4% in Extended Data Fig. 1b). Inset width, 100 nm. **e,** Archived astrocyte-associated protein-assay comparisons, expressed as log2(GLAST-positive sEV/total EV); n = 3 displayed assay records per marker, selected by the recorded effect-based rule in Methods; archived source records are listed in Supplementary Table S13. **f,** ApoB100 (lipoprotein) and platelet CD41–CD61 depletion-control ELISAs, expressed as log2(GLAST-positive sEV/GLAST-negative EV); n = 3 assay records per marker. ApoB100 was lower in all three records; CD41–CD61 was lower in two of three records and variable across records. **g,** RNA-biotype composition across participants; mRNA and lncRNA accounted for approximately 72% and 27% of exon-mapped reads, respectively, with similar group-level composition. **h,** Detected mRNAs, transcript isoforms, mature miRNAs and isomiRs; boxes, interquartile range; white lines, median; points, participants. Two-sided Mann–Whitney tests; n = 45 ALS and 15 controls. **i,** Sequence-motif atlas spanning miRNA EXOmotifs and mRNA 3′-UTR/RNA-binding-protein motifs. A total of 130 entries were retained, including 49 with an RNA-binding-protein assignment, defining an RBP-linked sequence context across both RNA classes. ALS–control prevalence was tested by two-sided Fisher exact tests with Benjamini–Hochberg correction; motif incidence was stable across groups after correction.

From each participant, a 20µl RNA eluate generated one small RNA and one transcriptome library, yielding 120 matched libraries across the 60-person discovery cohort (Fig. 1g,h). Exon-mapped reads were predominantly mRNA (approximately 72%) and long non-coding RNA (approximately 27%). Group level biotype composition and mature miRNA/isomiR richness were similar between ALS and controls. Transcript-isoform richness was lower in ALS (9,922 ± 6,058 versus 13,776 ± 6,146; nominal P = 0.036; Fig. 1h). Age, sex and sequencing depth contributed prominent variation in RNA composition and complexity, and gene-exon and transcript read depth correlated with observed survival (Extended Data Fig. 2a–l). These observations motivated explicit depth-sensitivity analyses of the survival-associated scores (Supplementary Note 2).

We next examined sequence features associated with RNA handling. The motif atlas contained 130 entries, including 49 with an RNA-binding protein (RBP) assignment, across miRNA EXOmotifs and mRNA 3′ untranslated-region motifs (Fig. 1i and Extended Data Fig. 3a–e). Y-box binding protein 1 (YBX1), heterogeneous nuclear ribonucleoproteins (hnRNPs) and related RNA-processing factors provide established links between sequence recognition and EV-RNA sorting.^40,42,49–52^ Motif-family comparisons showed no ALS-control difference after multiple-testing correction, placing these sequence features in the broader composition of the recovered compartment. A separate splice-junction audit detected 110 recurrent unannotated junctions with canonical splice-site motifs across the 60 profiles; the targeted screen recovered no qualifying published transactive response DNA-binding protein 43 (TDP-43)-dependent cryptic junctions at the obtained depth (Extended Data Fig. 4).

### Reciprocal miRNA–mRNA states organize the survival-associated program

The paired profiles provided the basis for identifying coordinated RNA variation associated with survival. We therefore used outcome-guided factor analysis to prioritize this variation within the molecular background. Multi-Omics Factor Analysis (MOFA+)^53,54^ was fitted in the 45 participants with ALS using 187 miRNAs, the 374 most variable mRNAs and survival as three views (Fig. 2a). Factor 1 accounted for 0.18% of feature-weighted molecular variance, compared with 8.2% and 13.1% for Factors 2 and 3. The inclusion of survival made Factor 1 a supervised prioritization axis within a much larger molecular background. In the complementary two-view model fitted without survival, no factor–survival association passed multiplicity correction (Supplementary Note 3). Candidate nomination therefore proceeded from the guided model.

**Fig. 2.**
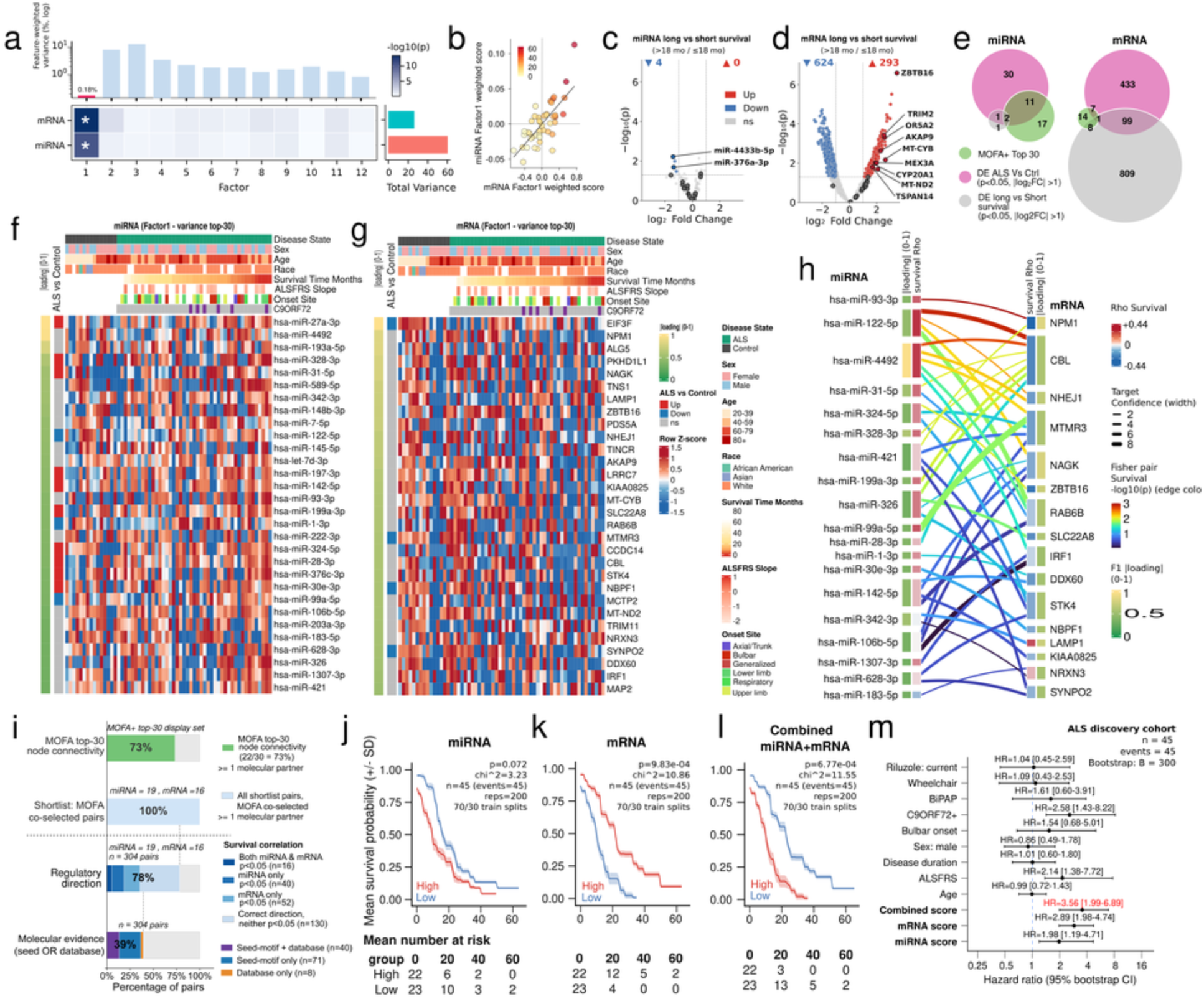
| Reciprocal miRNA–mRNA states reveal a low-variance survival program. Unless otherwise stated, analyses use the 45 ALS discovery participants. **a**, MOFA+ decomposition across miRNA, mRNA and survival views. Bars show feature-weighted variance explained across the two molecular views; the heatmap shows survival association. Factor 1 explained 0.18% of molecular variance, compared with 8.2% and 13.1% for Factors 2 and 3. Because survival was included as a model view, Factor 1 is outcome-guided by construction. **b**, Loading-weighted miRNA versus mRNA Factor 1 scores (Pearson r = 0.738, two-sided P = 7.07 × 10−9). None of 10,000 participant-identity permutations reproduced the observed cross-layer correlation (empirical P = 1/10,001). **c,d**, Differential abundance between long (>18 months) and short (≤18 months) survival for miRNAs and mRNAs; dashed lines mark nominal P = 0.05 and |log2 fold change| = 1 display thresholds. **e**, Overlap of Factor 1 top-loading, ALS–control and survival-associated display sets. The top-30 sets are display subsets only; candidate nomination used the prespecified top-50 loading prefilter. **f,g**, Row-scaled abundance of 30 high-loading miRNAs and 30 mRNAs across all 60 discovery profiles with clinical annotations. **h**, Highest-confidence target-supported links among the 19 nominated miRNAs and 16 mRNAs; ribbon width, composite target support; edge colour, Fisher-combined feature–survival statistic (−log10 P); side bars, feature loading and survival correlation. **i**, Evidence composition across the complete 19 × 16 cross-product: 238 of 304 pairs had opposing feature–survival directions and 119 contained a canonical seed match, a qualifying target-support record or both. **j–l**, Repeated-split Kaplan–Meier summaries for the miRNA, mRNA and combined scores across 200 stratified 70:30 splits. Weights and median cut points were estimated in 31-participant training subsets and applied to all 45 participants. These are full-cohort resubstitution summaries. Lines, mean survival; bands, split-to-split variability; displayed two-sided log-rank P and χ², medians across 200 tests. **m**, Separate univariable full-cohort Cox estimates for standardized molecular and clinical predictors; points, hazard ratios; bars, participant-bootstrap 95% intervals from 300 resamples. Row-specific n is reported in Source Data and Supplementary Table S14. ALSFRS-R decline was reverse-coded so that larger values denote faster decline.

Loading-weighted miRNA and mRNA Factor 1 scores correlated across participants (Pearson r = 0.738, P = 7.07 × 10⁻⁹; Fig. 2b). None of 10,000 participant-identity permutations reproduced this covariance (empirical P = 1/10,001; Supplementary Note 4), supporting the contribution of the matched design. Clinical overlays further describe the paired scores in participants with available ALSFRS-R and progression-slope data (Extended Data Fig. 5a,b). Integration of factor loading, differential-abundance contrasts and survival associations nominated 19 miRNAs and 16 mRNAs for subsequent analyses (Fig. 2c–g and Extended Data Fig. 5c–g; Supplementary Note 5). Supplementary Table S1 summarizes their disease-contrast and survival-stratum fold changes, with nominal P values and Benjamini–Hochberg (BH)-adjusted false discovery rates (FDRs), alongside the five miRNAs evaluated in total plasma. Supplementary Table S2 lists the four miRNAs that met the nominal display threshold without reaching FDR significance.

To determine how the cross-layer covariance was organized, we examined directional relationships among the nominated candidates. The 19 miRNAs and 16 mRNAs form 304 possible pairs, of which 119 carried a canonical TargetScan seed match, an experimental target record, or both.^45,55,56^ Across the full cross-product, 238 pairs (78.3%) had opposing feature– survival directions (Fig. 2h,i and Extended Data Fig. 6a–c,f; Supplementary Note 6). The miRNA-high/mRNA-low state accounted for 47.9% of prior-supported pairs in the shortest-survival tertile, compared with 25.5% and 21.9% in the intermediate– and longest-survival tertiles (short versus long, P = 0.012). Participant-level burden of this state correlated inversely with survival (r = −0.467, P = 0.0012; Extended Data Fig. 6d,e). Both reciprocal states separated survival, while the same-direction states yielded P = 0.621 and P = 0.685 (Extended Data Fig. 6g,h). Survival association thus depended on the relative configuration of the two RNA layers.

Combining the RNA layers increased apparent survival separation. Across 200 stratified 70:30 splits, weights estimated in the training subset and applied to the full cohort yielded median log-rank P values of 0.072 for miRNA, 9.8 × 10⁻⁴ for mRNA and 6.8 × 10⁻⁴ for the combined score (Fig. 2j–l). These curves summarize repeated full-cohort resubstitution. Separate standardized univariable Cox models fitted to the full discovery cohort gave hazard ratios of 1.98, 2.89 and 3.56, respectively (Fig. 2m). Both mRNA and combined scores retained associations after adjustment for gene-exon read depth: hazard ratio 3.13 (95% confidence interval (CI) 1.76–5.59; P = 1.10 × 10⁻⁴) for mRNA and 2.86 (95% CI 1.71–4.78; P = 6.00 × 10⁻⁵) for the combined score (n = 45). We next evaluated the miRNA candidates in an independent plasma cohort.

### Source-restricted miRNA discovery transfers to total plasma

To test whether the nominated program remained accessible outside the enriched fraction, we examined total plasma. All 19 sEV-nominated miRNAs were measurable in the Magen et al. cohort,^31^ within a set of 296 miRNAs shared between compartments (Fig. 3a and Extended Data Fig. 7a; Supplementary Note 7). Nine of the 19 candidates (47.4%) were differentially abundant between ALS and controls in plasma (Extended Data Fig. 7b–d). The corresponding proportions were 187 of 1,848 plasma-detected miRNAs (10.1%), 187 of 2,632 annotated miRNAs (7.1%) and 82 of 324 sEV-detected miRNAs (25.3%). Thus, the nominated set contained approximately 4.7 times the proportion of ALS-associated features found in the detected plasma background. Because transfer to total plasma is only informative if the prognostic relationship is retained, we next prioritized candidates whose survival associations preserved both effect magnitude and direction across compartments. Combining bootstrap Cox-effect magnitude with direction concordance between GLAST-positive sEVs and total plasma selected miR-31-5p, miR-4492, miR-93-3p, miR-421 and miR-324-5p, each with 97–100% cross-compartment direction concordance (Fig. 3b,c; Supplementary Tables S3 and S4; Supplementary Note 8). All five were also more abundant in ALS than controls in plasma (ratios 1.49–2.22; all BH FDR < 0.01; Supplementary Table S1). miR-1-3p showed no bootstrap direction concordance and was excluded; the original plasma study had likewise excluded it from prognostic testing due to longitudinal variability.

**Fig. 3.**
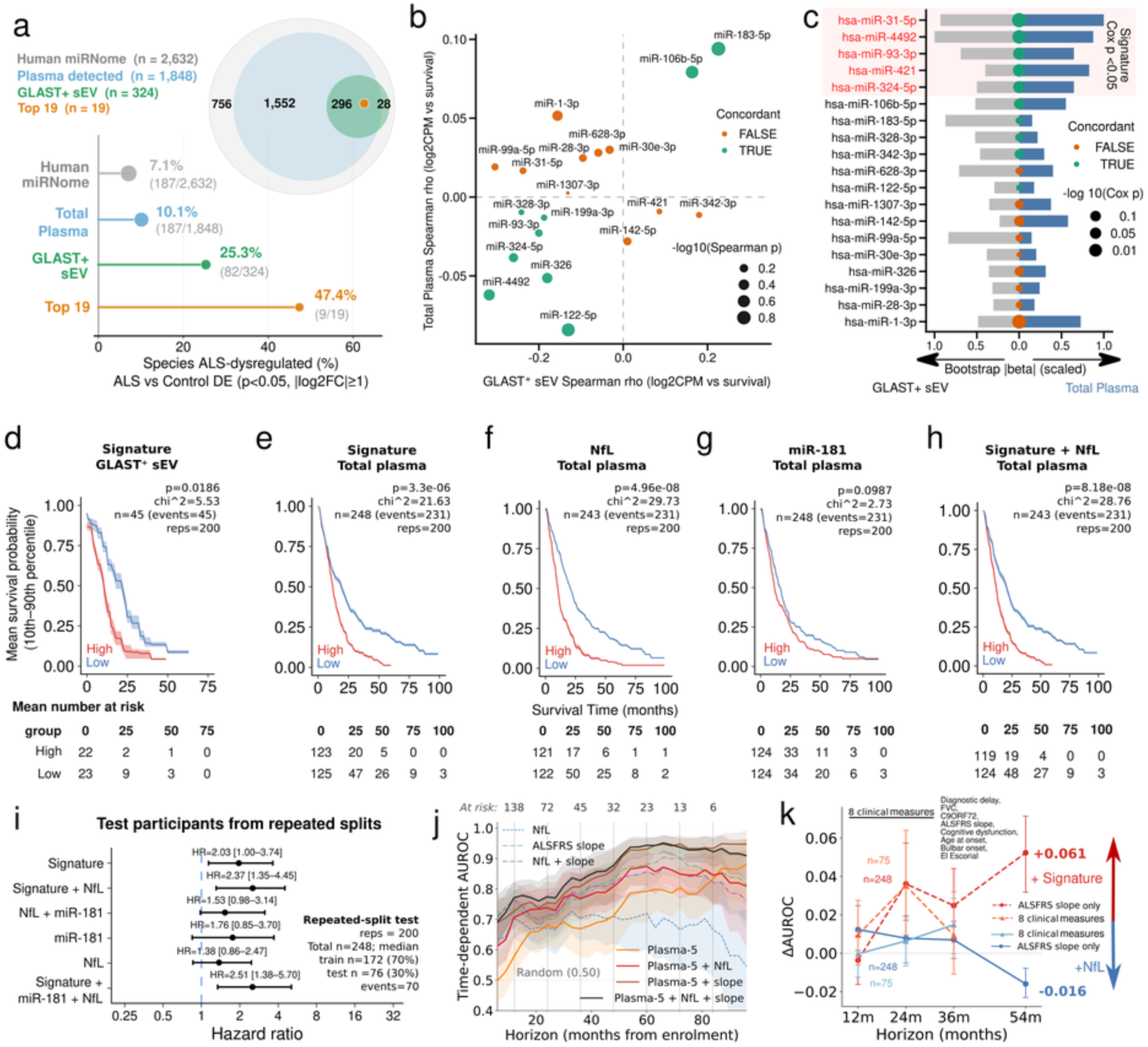
| Source-restricted miRNA discovery transfers to total plasma and adds prognostic information alongside NfL and ALSFRS-R. **a**, Cross-compartment coverage in GLAST-positive sEVs and the Magen et al.^31^ total-plasma cohort (248 ALS for survival, 231 deaths; 243 with NfL; 103 controls). Of 2,632 annotated miRNAs, 1,848 were detected in plasma, 324 in sEVs and 296 in both. Before plasma survival effects entered ranking, 9 of 19 sEV-nominated candidates were ALS–control differential in plasma (47.4%), compared with 187 of 2,632 annotated miRNAs (7.1%), 187 of 1,848 plasma-detected miRNAs (10.1%) and 82 of 324 GLAST-positive-sEV-detected miRNAs (25.3%). **b,** Candidate-wise Spearman survival correlations across compartments; colour indicates raw Spearman-sign agreement and point size plasma significance. **c,** Bootstrap ranking of all 19 candidates using scaled median absolute bootstrap Cox coefficients in GLAST-positive sEVs and plasma multiplied by cross-compartment direction concordance. The selected five are miR-31-5p, miR-4492, miR-93-3p, miR-421 and miR-324-5p. **d,** Repeated training-split/resubstitution Kaplan–Meier summary for the five-miRNA score in GLAST-positive sEVs (n = 45, 45 events; 200 stratified 70:30 splits); training-derived scores and cut-points were applied to all participants in each repeat, and displayed log-rank χ² = 5.53 and P = 0.0186 are medians across the 200 full-cohort tests. **e–h,** Corresponding repeated training-split/resubstitution summaries in total plasma for the five-miRNA score (n = 248), NfL (n = 243 with NfL available), miR-181 (n = 248) and five-miRNA score plus NfL (n = 243); curves show full-cohort resubstitution means with 10th–90th percentile ribbons. **i,** Hazard ratios evaluated in held-out test participants across the same splits; bars show the 2.5th–97.5th percentiles of split-specific adjusted estimates, with complete-case sample sizes varying for NfL-containing models. **j,** Horizon-specific full-cohort AUROC for the five-miRNA score and combinations with NfL and ALSFRS-R slope; participants censored before each horizon are excluded and bootstrap bands resample fixed scores without refitting the Cox model. **k,** Incremental cross-validated AUROC for RNA or NfL added to ALSFRS-R slope and the eight-variable clinical baseline; prediction-row bootstrap intervals are not participant-clustered. The incremental contribution of the five-miRNA score was greater than that of NfL at 24, 36 and 54 months over the ALSFRS-R slope baseline.

The five-miRNA score separated survival in repeated full-cohort resubstitution analyses in both discovery (median log-rank χ² = 5.53, P = 0.0186; n = 45; Fig. 3d) and plasma (median P = 3.3 × 10⁻⁶; Fig. 3e), and correlated with observed survival time in both compartments (Spearman ρ = −0.56 and −0.22, respectively; Extended Data Fig. 7e,f). NfL showed stronger full-cohort separation (P = 5.0 × 10⁻⁸; Fig. 3f), while miR-181 served as the published RNA comparator (Fig. 3g). In held-out analyses, median hazard ratios were 2.03 for RNA and 2.37 for RNA combined with NfL (Fig. 3h,i). An outcome-permutation analysis that repeated univariable Cox ranking within the 19-candidate set yielded a C-index of 0.5740 versus a null median of 0.5435 (empirical P = 0.026; Supplementary Note 4). Interparticipant variability and serial measurements further characterized the individual plasma features (Extended Data Fig. 7g–i,m; Supplementary Note 9).

The contribution of RNA varied with prediction horizon. In full-cohort analyses, RNA had higher area under the receiver operating characteristic curve (AUROC) than NfL at 24, 36 and 54 months, including 0.840 versus 0.704 at 54 months (Fig. 3j). In the separate cross-validated incremental analyses, adding RNA to ALSFRS-R slope changed AUROC by −0.013, +0.036, +0.039 and +0.061 at 12, 24, 36 and 54 months; corresponding NfL increments were +0.012, +0.007, +0.007 and −0.016 (Fig. 3k; Supplementary Table S5). An eight-variable clinical baseline in 75 complete cases provided a further comparison at 12, 24 and 36 months. At 24 months, RNA increased AUROC by 0.033 and NfL by 0.006; at 36 months, their increments were similar (0.014 and 0.015; Supplementary Table S6). Seven-classifier analyses describe variation across modeling choices (Extended Data Fig. 8a–i; Supplementary Note 10). The relative contribution of the two biomarkers was horizon dependent, with NfL contributing more at 12 months and the five-miRNA score providing the larger incremental gain at 24–54 months.

Processing batch was an important source of sensitivity in this retrospective plasma cohort. Batch covaried with survival (Kruskal–Wallis H = 61.3, P = 0.0009; Extended Data Fig. 7j). Empirical-Bayes ComBat correction^57^ changed the survival-correlation sign for 9 of the 19 candidates and attenuated repeated-split score separation to P = 0.125 (Extended Data Fig. 7k,l). The aligned technical and outcome structure makes the contributions of processing and biological sampling difficult to separate. The normalized-expression and ComBat analyses are reported together to show this sensitivity.

### Blood-nominated mRNAs show concordance with disease-relevant cortical programs

To provide biological context for the survival-associated paired-RNA program, we turned to the mRNA arm: the 16 candidates nominated in blood were evaluated in the New York Genome Center (NYGC) ALS Consortium resource analyzed by Eshima et al.^24^ (586 profiles from 308 donors) and the ALS/frontotemporal lobar degeneration (FTLD) single-nucleus resource of Pineda et al.^25^ (527,261 analyzed nuclei from 69 donors; Fig. 4a). This tissue bridge was important because a prognostic signal identified in circulation is more biologically compelling if its effector component is also represented in disease-relevant tissue programs rather than reflecting biofluid-specific variation alone. Disease-, region– and cell-class-resolved contrasts identified recurrent cortical differential expression among these candidates (Fig. 4b and Extended Data Fig. 9a,b). Their breadth of support varied by gene and contrast, motivating integration with the original sEV survival evidence as a tissue-context criterion. Classification using the resulting five-gene core further described separation from control and FTLD comparators across tissue contexts (Extended Data Fig. 9c–e). The classification analyses used the same tissue resources that informed core selection.

**Fig. 4.**
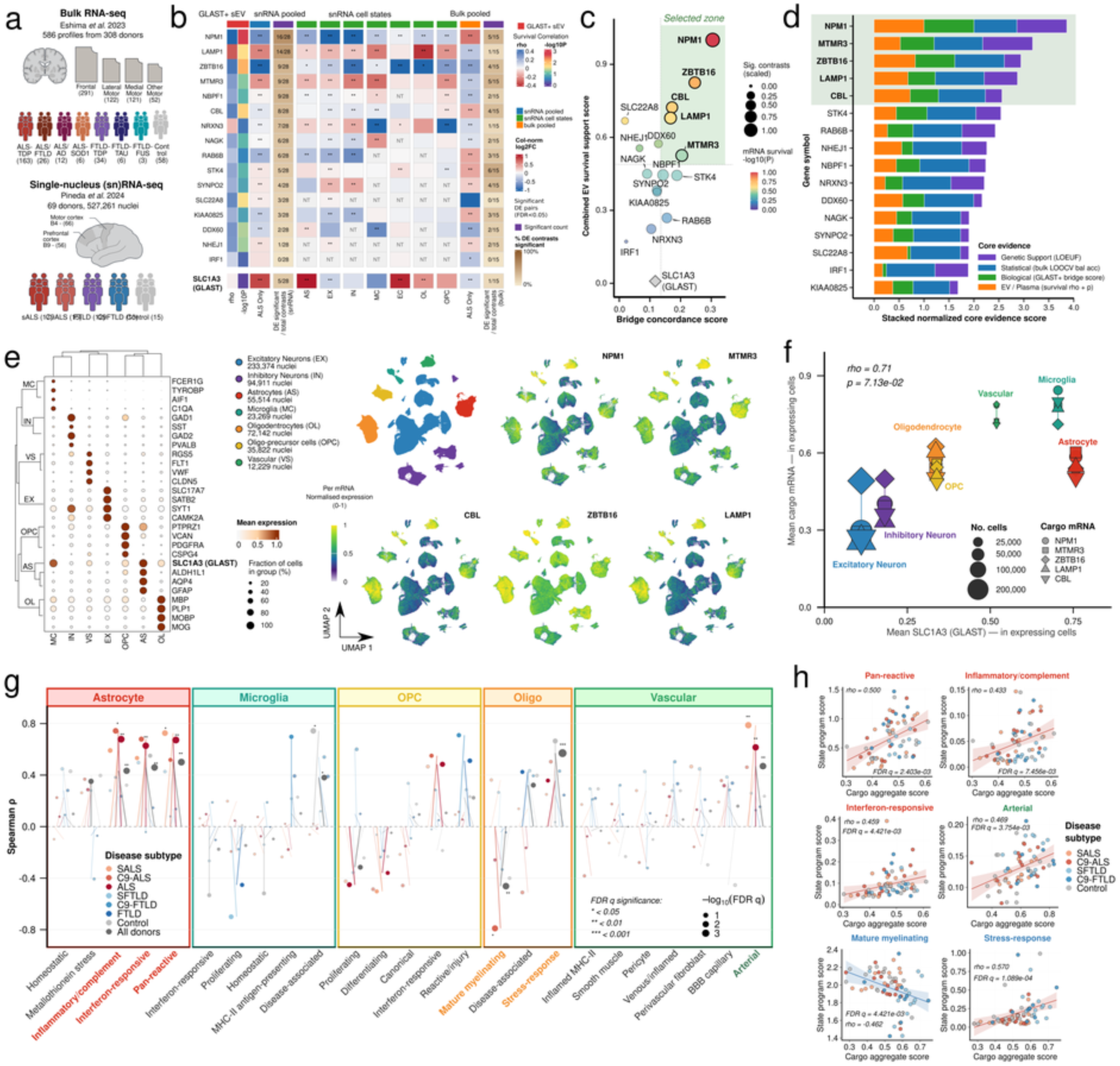
| Blood-nominated mRNAs show concordance with reactive glial–vascular programs in human ALS cortex. **a**, Independent cortical resources used to evaluate the 16 sEV mRNA candidates, fixed before tissue analysis: Eshima et al.^24^ bulk RNA-seq (586 profiles from 308 donors) and Pineda et al.^25^ single-nucleus cortex (527,261 analyzed nuclei from 69 donors across BA4 and BA9). Donor is the inferential unit for single-nucleus analyses. SLC1A3 is shown separately as the GLAST capture/reference marker. **b,** Cross-resource evidence matrix for the 16 candidate mRNAs. Fill, log2 fold change; asterisks, FDR-significant differential expression; NT, contrast not tested or not estimable. **c,** Candidate selection on tissue-bridge concordance and sEV survival support. Dashed lines mark candidate medians; the upper-right conjunction selects NPM1, MTMR3, ZBTB16, LAMP1 and CBL. **d,** Four-component sensitivity ranking combining gene-constraint/variant-context annotation, bulk classification, tissue bridging and EV/plasma survival evidence; the top five in this contextual ranking match the primary conjunction. **e,** Canonical broad-cell markers, uniform manifold approximation and projection (UMAP) of analyzed nuclei and normalized expression of each selected cargo mRNA across cell classes; the selected genes are expressed across broad cortical lineages. **f,** Mean cargo-mRNA expression versus mean SLC1A3 among expressing cells across seven broad classes (Spearman ρ = 0.71, two-sided P = 0.071); the seven-class relationship is descriptive. **g,** Donor-pseudobulk association of the aggregate five-gene score with 25 prespecified lineage-gated state programs across astrocyte, microglial, OPC, oligodendrocyte and vascular compartments; point size, significance; asterisks, BH-adjusted q values. None of the five selected cargo genes is included in a state-program gene set. **h,** Representative all-donor relationships for astrocyte pan-reactive, inflammatory/complement and interferon-responsive programs, vascular arterial identity, mature myelinating oligodendrocytes and oligodendrocyte stress. Blood and cortex were profiled in separate cohorts.

Five genes met the joint sEV-survival and cortical-support criterion. NPM1, MTMR3, ZBTB16, LAMP1 and CBL exceeded the candidate median on both the sEV-survival and tissue-bridge axes, whereas SLC22A8 and STK4 were supported on a single axis (Fig. 4c; Supplementary Table S7). The convergence of survival support in circulating sEVs with recurrent cortical evidence was notable: the same five genes were also ranked highest by a separate four-component score incorporating bulk classification and gene-constraint/variant context. A separate four-component score incorporating bulk classification and gene-constraint/variant context ranked the same five genes highest (Fig. 4d; Supplementary Table S8). The core was expressed across major cortical lineages (Fig. 4e). Across seven broad cell classes, its mean expression showed a positive descriptive relationship with SLC1A3 expression (Spearman ρ = 0.71, P = 0.071; Fig. 4f). We therefore examined core-associated states within each lineage.

The aggregate cargo score correlated positively with astrocyte pan-reactive, inflammatory/complement and interferon-responsive programs, vascular arterial identity and oligodendrocyte stress, and inversely with mature myelinating oligodendrocyte identity (|ρ| = 0.433–0.570; all q < 0.008; Fig. 4g,h). Several associations were also present within ALS, linking the score to heterogeneity inside the diagnosis. At gene level, 43 cargo–program associations passed q < 0.05: 18 for LAMP1, 10 for ZBTB16, six each for NPM1 and MTMR3, and three for CBL (Extended Data Fig. 10b). Program gene sets excluded the five cargo genes, and shared response programs were evaluated within prespecified parent lineages (Extended Data Fig. 10a; Supplementary Note 11). These associations show that the selected core is represented within glial and vascular responses described in ALS cortex.^17–19,21–25^

### Paired RNA candidates converge on ALS-relevant glial–vascular pathways

We next asked whether the paired RNA candidates and the independent cortical programs implicated common biological pathways. A source-attributed network assembled with EVd3x^58^ connected the five selected miRNAs and five cortex-supported mRNAs through scored intermediate relationships and target evidence (Fig. 5a). Enrichment of lineage-gated cortical programs against Gene Ontology, Reactome and MSigDB Hallmark terms^59–61^ identified interferon signaling and antigen presentation, inflammatory/complement activity, oxidative stress, vascular activation and reduced myelination and axon–glia support (Fig. 5b). These themes align with the astrocytic,^17–19,23–25^ microglial,^20,25^ oligodendroglial^21,25^ and neurovascular^22,25^ responses represented in the cortical analyses.

**Fig. 5.**
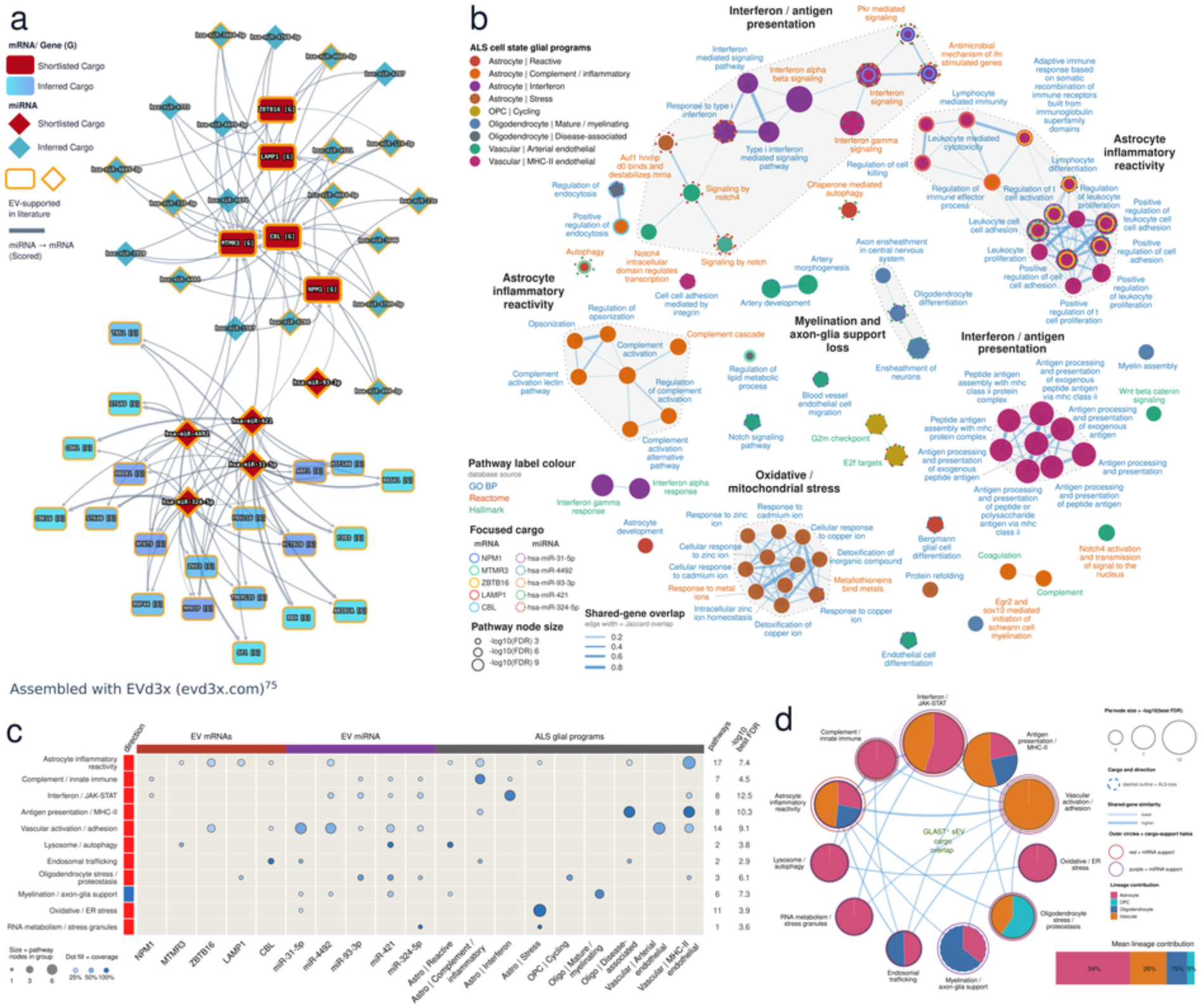
| Paired RNA candidates converge on ALS-relevant glial–vascular pathways. **a**, Evidence-synthesis network assembled with the source-attributed EVd3x platform (evd3x.com)^58^, connecting the five selected miRNAs and five cortex-supported mRNAs through scored intermediate relationships. Edges show scored miRNA–mRNA relationships; gold outlines mark literature-supported EV cargo. Red nodes identify shortlisted cargo and blue nodes inferred intermediates. **b,** Enrichment map of lineage-gated cortical programs against Gene Ontology, Reactome and MSigDB Hallmark terms. Node fill denotes the lineage/state program, label colour the pathway database, node size −log10 FDR and edge width shared-gene Jaccard overlap. Recurrent territories include interferon/antigen presentation, astrocyte inflammatory reactivity, oxidative/mitochondrial stress, vascular activation and loss of myelination/axon–glia support. **c,** Eleven modules receiving support from both RNA arms. Columns show selected mRNAs, predicted miRNA target footprints and lineage-gated cortical programs; point size indicates pathway count and fill coverage; the right-hand column reports best −log10 FDR. Ten modules increase in ALS, while myelination/axon–glia support decreases. **d,** Integrated module map. Pie sectors show normalized lineage enrichment contribution, and edges show shared support across modules. Mean lineage contribution is 54% astrocyte, 26% OPC, 15% oligodendrocyte and 5% vascular. The integrated view resolves interferon/JAK–STAT, antigen presentation, inflammatory/complement, vascular and lysosomal/endosomal programs together with loss of myelin and axon support.

The mRNA program and predicted miRNA target footprints overlapped 11 shared modules, ten associated with increases in ALS and one with reduced myelination/axon–glia support (Fig. 5c). Interferon/Janus kinase–signal transducer and activator of transcription (JAK–STAT) signaling had the strongest reported enrichment (eight pathways; best −log10 FDR, 12.5), followed by major histocompatibility complex class II (MHC-II) antigen presentation (eight pathways; 10.3) and vascular activation/adhesion (14 pathways; 9.1). Astrocyte inflammatory reactivity spanned the most pathways (17; 7.4). Astrocytes accounted for 54% of normalized pathway-enrichment contributions, followed by OPCs (26%), oligodendrocytes (15%) and vascular cells (5%; Fig. 5d). Lysosome/autophagy and endosomal trafficking formed another recurrent theme, connecting the LAMP1-containing core with cellular systems relevant to RNA handling and vesicle cargo composition.^40,42,49–51^ The shared modules provide pathway-level context linking inflammatory reactivity and altered trafficking with reduced myelin support.

## Discussion

Paired RNA profiling in GLAST-positive small extracellular vesicles isolated from plasma of ALS patients identified a survival-associated program organized across miRNA and mRNA layers. Reciprocal miRNA-high/mRNA-low configurations tracked shorter survival, while target-supported relationships provided a directional framework for the joint signal. The miRNA arm transferred to total plasma, whereas the mRNA arm showed concordance with glial, vascular and myelination-related programs in independent cortical datasets. The primary contribution is therefore a paired circulating-RNA strategy that identifies a low-variance prognostic program and assigns it biological context.

The sampling strategy was central to this result by retaining paired miRNA and mRNA measurements from the same surface-selected compartment in each participant. The survival-associated factor occupied only 0.18% of feature-weighted molecular variance, against a background dominated by demographic and technical variation. Outcome-guided integration brought this small component into focus, and all 19 nominated miRNAs were subsequently measurable in total plasma. Refinement to five candidates shows how discovery in an enriched fraction can inform assay development in a more accessible matrix. Measuring both RNA classes in the same preparations linked miRNA nomination to the mRNA program carried forward for tissue contextualization.

Within these paired measurements, the joint RNA state was more informative than either layer alone in discovery. Shorter survival was associated with greater miRNA-high/mRNA-low burden, while reciprocal configurations separated survival among the same candidates. Canonical seed matches and experimental target records^45,55,56^ place these relationships within an established post-transcriptional framework, but do not establish direct target engagement. RBP-associated motifs add a related observation: recovered RNAs contain sequences recognized by YBX1, hnRNPs and other factors involved in extracellular cargo sorting.^40,42,49–52^ Together, these data support a directionally organized regulatory interpretation that can be tested experimentally.

The paired mRNA arm supplied tissue-level context for the circulating program. The selected genes were recurrently dysregulated across independent cortical resources and their aggregate score associated with reactive astrocyte, oligodendrocyte stress, reduced myelinating identity and vascular programs.^17–19,21–25^ Several associations persisted within ALS, linking the core to variation among affected donors. The representation of the selected mRNAs across OPC, oligodendrocyte and vascular compartments, although less prominent than in astrocytic programs, is consistent with GLAST-positive EV enrichment rather than lineage specificity. This broader representation may help explain why the circulating program converged on coordinated glial–vascular state programs rather than an exclusively astrocytic response. Excluding cargo genes from the state-program definitions reduced direct mathematical overlap, and lineage gating localized shared stress and immune responses to their parent cell classes. These analyses support biological concordance between the circulating mRNA candidates and disease-relevant cortical programs.

The convergence of candidate and cortical pathway analyses on vesicle trafficking raises a specific mechanistic hypothesis for follow-up. LAMP1 belongs to the cortex-supported core, lysosomal and endosomal pathways recur in the integrated analysis, and the recovered cargo contains RBP-associated sorting motifs. One possibility is that reactive cellular programs alter both RNA abundance and extracellular release. Paired cellular and vesicle measurements during targeted perturbation could test these steps, including RNA packaging and target engagement. Longitudinal sampling would establish whether changes in the circulating program accompany changes in pathway activity or treatment response.

The prediction results place the RNA score alongside established prognostic measures and suggest why the circulating signal may be clinically useful. RNA contributed more discrimination at later horizons when added to functional decline, whereas NfL retained strong overall prognostic information. NfL reflects neuroaxonal injury,^7–13^ while the tissue contextualization indicates that the paired RNA candidates intersect glial, vascular and myelin-related biology, indicating nonredundant components of disease progression. The stronger contribution of the miRNA score at longer survival horizons is consistent with emerging evidence that NfL is particularly informative near major clinical transitions, whereas broader molecular signatures can provide complementary information over longer intervals.^9,15,16^ The present analysis extends circulating-miRNA prognostic work, including the miR-181 study,^31^ by linking candidate selection to a paired mRNA framework and independent tissue context. A fixed plasma score evaluated alongside NfL and clinical predictors would test whether this distinction improves prognosis and patient stratification. Horizon-specific biomarker information may be useful for prognostic enrichment or stratified randomization when trial duration and expected event timing differ.^16^

The study has several limitations. Survival guided discovery in a small cohort, and the plasma and cortical resources contributed to refinement of the final panels. Full-cohort survival displays therefore report apparent separation; generalizability requires prospective testing of a fixed assay and score. Plasma batch covaried with outcome, and ComBat showed substantial preprocessing sensitivity. GLAST capture enriches an astrocyte-associated circulating compartment but is not entirely astrocyte specific, as also suggested by the broader cortical distribution of the nominated mRNAs. The cortical analyses should therefore be interpreted as cross-cohort biological context, not evidence that circulating RNA measures an individual’s cortical state. Lineage tracing, single-vesicle localization and paired blood–tissue studies are needed to establish origin, co-packaging and individual correspondence.

Paired extracellular-vesicle RNA profiling identifies an ALS survival-associated miRNA–mRNA program within a source-enriched circulating compartment. Its miRNA arm transfers to total plasma, while its mRNA arm shows concordance with disease-relevant cortical programs involving glial reactivity and reduced myelin support. More broadly, this study provides a framework for multiview, multi-omic analysis within cell-type-enriched circulating compartments, where source enrichment can reduce biological heterogeneity while paired molecular layers preserve regulatory organization. Extending this approach across additional cell-type-enriched EV populations could enable coordinated analysis of astrocytic, oligodendroglial, vascular and other disease-associated programs in circulation. The ALS panels provide a tractable starting point for prospective evaluation of a fixed assay and score.

## Methods

### Study participants and design

The discovery cohort comprised 45 participants with ALS and 15 non-neurological controls, with paired small-RNA and messenger-RNA profiles generated from GLAST/SLC1A3-positive plasma sEVs. Three published resources extended the analysis: the Magen et al.^31^ total-plasma small-RNA cohort (Gene Expression Omnibus (GEO) GSE168714; 248 participants with ALS for survival analyses, 243 with NfL measurements and 103 controls for the case–control contrast), the New York Genome Center (NYGC) ALS Consortium bulk-cortex resource analyzed by Eshima et al.^24^ (GEO GSE153960; 586 profiles from 308 donors), and the Pineda et al.^25^ ALS/frontotemporal lobar degeneration (FTLD) single-nucleus resource (National Center for Biotechnology Information (NCBI) PRJNA1073234; Synapse syn51105515; 625,973 published nuclei from 73 donors, of which 527,261 nuclei from 69 donors had complete annotation for the analyses shown). Table 1 and Fig. 1a summarize cohort characteristics and analytical roles. Supplementary Table S9 lists per-variable availability and analytical role across the blood cohorts.

**Table 1.** | Characteristics of the discovery, plasma-transfer and cortical resources. Values are mean ± standard deviation (s.d.) unless otherwise indicated. Dashes denote unavailable or not applicable data. Donor counts are shown for cortical resources, with the corresponding numbers of sequenced profiles or analyzed nuclei listed under total analyzed units. Blood-cohort demographic and clinical summaries refer to ALS participants; control counts are listed separately. Discovery ages at collection are means of reported age-band midpoints. Cortical summaries retain the donor or profile denominators shown. The plasma cohort is from Magen et al.^31^, bulk cortex from Eshima et al.^24^ and single-nucleus cortex from Pineda et al.^25^. Revised ALS Functional Rating Scale (ALSFRS-R) decline is expressed as points lost per month in both blood cohorts.

| Characteristic | Blood cohorts |  | Cortical resources |  |
| --- | --- | --- | --- | --- |
|  | GLAST-positive sEV discovery | Total-plasma transfer and refinement (Magen et al.<sup>31</sup>) | Bulk cortical resource (Eshima et al.<sup>24</sup>) | Single-nucleus cortical resource (Pineda et al.<sup>25</sup>) |
| Cohort / source | National ALS Biorepository; present study | Magen et al.; GEO GSE168714 | NYGC ALS Consortium; GEO GSE153960 | Pineda et al.; PRJNA1073234; Synapse syn51105515 |
| Participants |  |  |  |  |
| ALS, n | 45 | 248 | 163 ALS-TDP; 26 ALS/FTLD; 12 ALS/AD; 6 ALS-SOD1 donors | 32 donors (17 sporadic, 15 C9orf72) |
| Control, n | 15 | 103 | 58 donors | 15 donors |
| Other disease, n | — | — | 34 FTLD-TDP; 6 FTLD-tau; 3 FTLD-FUS donors | 22 FTLD donors (12 sporadic, 10 C9orf72) |
| Total analyzed units, n | 60 participants | 248 ALS for survival; 351 for disease contrast | 586 profiles from 308 donors | 625,973 published nuclei from 73 donors; 527,261 nuclei from 69 donors analyzed here |
| Demographics |  |  |  |  |
| Age at sampling or death, years | 63.7 ± 8.7 | 65.1 ± 11.1 | 65.2 ± 9.8 | 70.3 ± 10.0 |
| Age at onset or diagnosis, years | 62.8 ± 9.8; n = 35 | 62.7 ± 11.4 | 61.5 ± 10.4 | 62.6 ± 7.1 (motor onset) |
| Female, n (%) | 24 (53) | 144 (58) | 267 profiles (46) | 37 donors (54) |
| Clinical characteristics |  |  |  |  |

|  | GLAST-positive<br>sEV discovery | Total-plasma<br>transfer and<br>refinement<br>(Magen et al.<sup>31</sup>) | Bulk cortical<br>resource<br>(Eshima et al.<sup>24</sup>) | Single-nucleus<br>cortical<br>resource<br>(Pineda et al.<sup>25</sup>) |
| --- | --- | --- | --- | --- |
| Cohort / source | National ALS<br>Biorepository;<br>present study | Magen et al.;<br>GEO<br>GSE168714 | NYGC ALS<br>Consortium;<br>GEO<br>GSE153960 | Pineda et al.;<br>PRJNA1073234;<br>Synapse<br>syn51105515 |
| ALSFRS-R total<br>score | 28.6 ± 11.3; n =<br>35 | 35.9 ± 7.8 | — | — |
| ALSFRS-R decline,<br>points lost per month | 0.7 ± 0.5; n = 18 | 0.7 ± 0.7 | — | — |
| Disease duration at<br>sampling, months,<br>median (IQR) | 34.2 (18.1–46.2);<br>n = 27 | — | — | — |
| C9orf72 expansion | 6 positive; 39<br>negative | 32 positive (13%) | — | 25 of 69 donors:<br>15 C9-ALS; 10<br>C9-FTLD |
| Tissue / region |  |  |  |  |
| Region | Peripheral blood | Peripheral blood | Frontal 291;<br>lateral motor<br>122; medial<br>motor 121; other<br>motor 52 profiles | BA4, 66 donors;<br>BA9, 56 donors |
| Molecular subtype | — | — | OX 239; TE 128;<br>GLIA 84; control<br>135 profiles | Excitatory,<br>inhibitory, glial<br>and vascular<br>classes |
| Outcome |  |  |  |  |
| Deaths, n (%) | 45 (100) | 231 (93) | All post-mortem | All post-mortem |
| Survival from<br>collection, months,<br>median (IQR) | 13.0 (7.0–24.0) | — | — | — |
| NfL available, n (%) | — | 243 (98) | — | — |

ALS plasma was obtained from the National ALS Biorepository, administered by the Agency for Toxic Substances and Disease Registry (ATSDR) within the US Centers for Disease Control and Prevention (CDC) National ALS Registry.^62^ Specimens and linked clinical variables were provided as coded materials. Participants enrolled in the National ALS Registry provided written informed consent for biospecimen collection and research use under the National ALS Biorepository protocol, and secondary analysis of the coded specimens was approved by the Yale University Institutional Review Board (protocol 2000031350). Available variables included age, sex, site of onset, disease duration, ALSFRS-R, progression slope, C9orf72 status, riluzole use and respiratory/supportive-care variables. Survival was measured from enrollment to death or censoring, and survival time remained paired with censoring status in every permutation. Control plasma was collected under Yale University Institutional Review Board protocol 2000031350 with written informed consent from every control participant. Controls were used for compartment characterization and ALS-versus-control differential-abundance analyses; MOFA+ fitting, directional-state analyses and survival modeling were restricted to participants with ALS.

### Plasma processing and GLAST-positive sEV isolation

Whole blood was collected in dipotassium ethylenediaminetetraacetic acid (K2EDTA) tubes, gently inverted 5–10 times and processed within 2 h. Plasma was separated at 2,000 × g for 15 min at room temperature, transferred without disturbing the buffy coat and centrifuged at 3,500 × g for 10 min to obtain platelet-poor plasma. Aliquots were stored at −80 °C without repeated freeze–thaw cycles before isolation.

GLAST-positive sEVs were isolated from 0.5 ml platelet-poor plasma by total-EV precipitation followed by streptavidin–biotin immunocapture. Plasma received 10 µl Halt Protease and Phosphatase Inhibitor Cocktail (100×; Thermo Scientific, 1861282), was briefly cleared at 12,000 × g, and was then centrifuged at 1,500 × g for 15 min and 14,000 × g for 40 min at 4 °C. Fibrinogen was removed with 7.5 µl thrombin (System Biosciences, TMEXO-1) for 30 min at room temperature followed by 10,000 × g for 5 min at 4 °C. Total EVs were precipitated with ExoQuick (125 µl per 0.5 ml plasma; System Biosciences, EXOQ20A-1) and recovered at 2,500 × g for 30 min at 4 °C, followed by a second 2,000 × g spin for 5 min to consolidate the pellet.

In parallel, 50 µl Pierce Streptavidin Plus UltraLink resin (Thermo Scientific, 53117) per sample was conjugated with 8 µl biotinylated anti-GLAST/SLC1A3 (ACSA-1; Miltenyi Biotec, 130-118-984) and washed to remove unbound antibody. Total EVs were resuspended in 400 µl phosphate-buffered saline (PBS) containing inhibitor; 15 µl was retained for nanoparticle tracking and the remainder was incubated with anti-GLAST resin overnight at 4 °C with rotation. Bead-bound sEVs were collected at 400 × g for 10 min at 4 °C, washed twice with 1 ml PBS and stored at −80 °C. GLAST enrichment defines a surface-selected, glial-enriched discovery compartment; sEV is used operationally according to the measured size range and MISEV2023 terminology.^48^

### sEV characterization

GLAST-positive preparations were characterized by nanoparticle tracking analysis (NTA), immunogold transmission electron microscopy (TEM), three-colour direct stochastic optical reconstruction microscopy (dSTORM), nano-flow cytometry and protein assays (Fig. 1b–f; Extended Data Fig. 1). The platforms provide complementary measurements of particle size and concentration, ultrastructure, tetraspanin phenotype and biochemical enrichment. EV-method reporting is summarized in Supplementary Tables S10 and S11, with reagent records in Supplementary Table S12.

### Nanoparticle tracking analysis

Particle size and concentration were measured on a ZetaView x30 instrument (Particle Metrix) with a 405-nm laser using ZetaView v8.04.04. The instrument was calibrated with polystyrene standards before acquisition. Samples were recorded at 11 positions at 25 °C with a 5-nm bin width; analysis used minimum brightness 25, particle area 5– 1,000 and minimum trace length 15. The GLAST-positive fraction was calculated as captured-particle concentration divided by the matched total-particle concentration from the same plasma. Ten technical characterization preparations, separate from the 60 sequenced participants, are shown in Fig. 1b.

### Immunogold TEM

GLAST was detected with biotinylated anti-GLAST and 10-nm streptavidin-gold (Abcam, ab270041). Four microlitres of gold conjugate was incubated with 50 µl EV suspension overnight at 4 °C. A 4-µl aliquot was adsorbed for 2 min to a glow-discharged 200-mesh carbon-coated copper grid, washed, negatively stained with 2% uranyl acetate for 2 min and imaged on an FEI Tecnai transmission electron microscope at 80 kV with an AMT NanoSprint 15 MKII camera.

### dSTORM

Single-particle tetraspanin imaging was performed on a Nanoimager S Mark II (Oxford Nanoimaging; 100×, 1.4-numerical-aperture objective) using the EV Profiler 2 assay. GLAST-positive particles were captured with biotinylated anti-GLAST, incubated on the capture surface for 75 min and detected with antibodies to CD9, CD63 and CD81 for 50 min after a 15-min surface/capture preparation. The 488-, 561– and 640-nm channels were acquired sequentially in total internal reflection fluorescence (TIRF) mode. Localizations were filtered in NimOS v1.18.3 and clustered in ONI CODI using density-based spatial clustering of applications with noise (DBSCAN; 85-nm neighborhood); six fields of view were acquired per lane. Phenotype proportions were calculated after localization and clustering.

### Nano-flow cytometry

Samples were acquired on a CytoFLEX Nano (Beckman Coulter) after daily quality control, filtered-water background confirmation (≤100 events s−1) and nanoscale sizing calibration. Tetraspanins were detected with fluorescein isothiocyanate (FITC)-anti-CD9 (BD Biosciences, 555371), allophycocyanin (APC)-anti-CD63 (Immunotech, A87789) and APC-anti-CD81 (Immunotech, IM1165U). Antibody aggregates were removed at 20,000 × g for 10 min. EVs were initially diluted 1:25, and 20 µl EV suspension was incubated with 20 µl antibody mixture for at least 1 h at room temperature in the dark before further dilution. Violet side scatter served as the particle trigger; unstained and single-stain controls defined fluorescence gates, and acquisition was maintained below 2,000 events s−1 to reduce swarm detection. Data were acquired in CytExpert Nano v1.2.0.34, calibrated with FCM PASS v5.0.11^63^ and analyzed in FlowJo v10.10.

### Protein assays

GFAP (EMD Millipore/Sigma, NS830), AQP4 (Cusabio, CSB-E08254h), ALDOC (LSBio, LS-F7159) and GLT-1/SLC1A2 (LSBio, LS-F8708) were measured by commercial sandwich enzyme-linked immunosorbent assays (ELISAs) after elution of bead-bound EV material in glycine-HCl and neutralization. Enrichment was calculated as log2(GLAST-positive sEV/total EV). Apolipoprotein B100 (ApoB100; LSBio, LS-F22513-1; archived assay v2.0, 20 February 2026) and platelet CD41–CD61 (LSBio, LS-F56371-1; archived assay v1.0, 27 February 2026) were measured as depletion controls in separate commercial sandwich ELISAs and expressed as log2(GLAST-positive sEV/GLAST-negative EV). For the recorded depletion-control assays, bead-bound EVs were washed, eluted in 250 µl 0.1 M glycine-HCl (pH 2.5) and neutralized with 35 µl 1 M Tris-HCl (pH 8.0) before plating. Standards, samples or blanks (100 µl per well) were incubated for 90 min at 37 °C, followed by 100 µl biotinylated detection antibody for 60 min at 37 °C. ApoB100 plates were not washed before detection; platelet plates were washed twice at this step. After three washes, 100 µl horseradish peroxidase (HRP)–streptavidin was incubated for 30 min at 37 °C, followed by five washes, 90 µl 3,3′,5,5′-tetramethylbenzidine (TMB) substrate for 15 min (ApoB100) or 10–20 min (platelet), 50 µl stop solution and immediate absorbance measurement at 450 nm. Assay-specific sample dilutions followed the recorded plate maps. Depletion-control ratios were based on assay concentration without particle-count or total-protein normalization. Cargo-marker comparisons are retained as archived assay records in Supplementary Table S13. In the supplied Figure 1e record selection, up to three records per marker were ranked by |log2 ratio| + 0.03 × (total_n + glast_n); these records are descriptive and retain their assay-specific denominator.

### RNA extraction and sequencing

Total RNA was extracted from bead-bound GLAST-positive sEVs using the miRNeasy Serum/Plasma Advanced Kit (QIAGEN, 217204) without carrier RNA. Beads were suspended in 60 µl Buffer RPL plus 200 µl 1× Tris-buffered saline (TBS) for 3 min, pelleted at 400 × g for 10 min and the lysate was treated with 20 µl Buffer RPP for 3 min. After clarification at 12,000 × g for 3 min, an equal volume of isopropanol was added and the sample was loaded onto an RNeasy UCP MinElute column. Columns were washed with 700 µl RWT, 500 µl RPE and 500 µl 80% ethanol, dried at 12,000 × g for 5 min and eluted in 20 µl RNase-free water after a 10-min incubation. RNA was stored at −80 °C until library preparation.

### Small-RNA libraries

Libraries were prepared from 5 µl RNA eluate using the QIAseq miRNA Library Kit (QIAGEN). 3′ and 5′ adapters were ligated sequentially before complementary DNA (cDNA) synthesis and 12-nt unique-molecular-identifier (UMI) assignment, amplification and cleanup. Libraries were assessed on an Agilent TapeStation, size-selected when adapter dimer was excessive and quantified by KAPA quantitative polymerase chain reaction (qPCR) before pooling.

### Messenger-RNA libraries

Transcriptome libraries were prepared with the SMARTer Stranded Total RNA-Seq Kit v3 – Pico Input Mammalian (Takara) using 250 pg–10 ng total RNA. Ribosomal cDNA was depleted with ZapR v3 and mammalian R-probes, followed by bead purification, qPCR quantification and insert-size assessment. Libraries meeting the sequencing input criterion (≥0.5 ng µl−1 after final cleanup) were pooled.

### Sequencing

Libraries were normalized to 2.0 nM and sequenced at the Yale Center for Genome Analysis on an Illumina NovaSeq platform. Small-RNA libraries targeted 5–10 million passing-filter clusters per sample and transcriptome libraries approximately 25 million; sequencing was 100-bp paired-end with a 10-bp unique dual index. PhiX was included at 0.3% per lane as a run-quality control, and demultiplexing used CASAVA v1.8.2. The discovery dataset comprised 60 participants, 60 small-RNA FASTQ files and 120 paired-end transcriptome FASTQ files, corresponding to one small-RNA and one transcriptome library per participant.

### RNA quantification and preprocessing

Small-RNA reads were processed with the QIAseq miRNA GeneGlobe workflow. The 3′ adapter and common sequence were trimmed, 12-nt UMIs were extracted, reads sharing a UMI were collapsed to one molecule and deduplicated sequences were mapped to miRBase mature and hairpin references before genomic alignment. Per-sequence output retained isomiRs separately. Sequences mapping ambiguously to more than one miRBase entry were retained for richness summaries but excluded from feature-level modeling.

mRNA counts were generated from the paired transcriptome libraries. Raw counts, trimmed mean of M-values (TMM)-normalized log2 counts per million (CPM), variance-stabilized expression, ComBat-adjusted expression and MOFA inputs were retained as separate layers. For MOFA eligibility, the reported detection rule was log2CPM >5 in at least 10% of samples within a group; the mRNA view then retained the 374 most variable features. The analysis used edgeR/limma, DESeq2 and ComBat workflows.^57,64–66^ Supplementary Table S14 records the matrix and statistical test assigned to each panel, and Supplementary Note 2 describes batch and depth sensitivity. Supplementary Table S1 consolidates the additional disease and survival-stratum differential-abundance records. Ratios are oriented as ALS/control and ≥18/<18 months from collection; log2 fold changes are calculated as log2(ratio), with nominal P values and source-supplied BH FDRs retained separately.

### RNA composition, sequence motifs and splice junctions

RNA biotypes and feature richness were summarized from mapped annotations (Fig. 1g,h; Extended Data Fig. 2). ALS and control groups were compared by two-sided Mann–Whitney tests. Age-group displays used Kruskal–Wallis tests followed by Dunn pairwise comparisons with Holm adjustment, as specified in Extended Data Fig. 2c–h. Continuous clinical variables used Spearman correlation, and categorical variables used Mann–Whitney or Kruskal–Wallis tests as appropriate, with Benjamini–Hochberg (BH) correction within each analysis family.

Sequence motifs associated with extracellular-RNA sorting were evaluated in a universe of 162 eligible miRNAs and 6,097 protein-coding mRNAs with representative 3′ untranslated regions (UTRs) (Fig. 1i; Extended Data Fig. 3). The atlas included reported miRNA EXOmotifs, ATtRACT RNA-binding protein (RBP) motifs and literature-supported 3′-UTR motifs.^40,49,51^ ALS–control prevalence differences were tested by two-sided Fisher exact tests with BH correction within motif families. Supplementary Table S15 summarizes motif-family results; complete panel-level values are provided in Source Data Figure 1, sheet I.

Unannotated splice junctions were evaluated from read 1 after removal of the Illumina adapter sequence AGATCGGAAGAGCACACGTCTGAACTCCAGTCA with cutadapt v5.2^67^ (minimum retained length, 30 nt) and alignment to Genome Reference Consortium Human Build 38 (GRCh38)/GENCODE v50 with STAR v2.7.11 in two-pass mode.^68^ Junction-supporting molecules required a primary unique alignment, mapping quality (MAPQ) ≥30, zero mismatches and ≥15-nt anchors on both sides and were collapsed by UMI. Published KCNQ2, STMN2 and UNC13A TDP-43-associated junctions were queried at their reported coordinates.^69–74^ De novo discovery used first-pass STAR junctions only and required canonical splice motifs, ≥12-nt overhang and recurrence across participants. Junction burden was normalized to uniquely mapped depth before clinical association testing (Extended Data Fig. 4).

### Guided MOFA+ integration and candidate selection

Multi-Omics Factor Analysis (MOFA+)^53^ was fitted in the 45 participants with ALS using three views: 187 miRNAs, the 374 most variable mRNAs and survival. Molecular features were standardized within view and 12 factors were retained. The fitted object, feature filters, scaling parameters, convergence settings and random seed were archived with the analysis code. Factor-level molecular variance was calculated from the two RNA views only, excluding survival from the denominator. Survival was included deliberately to prioritize clinically linked structure, making Factor 1 an outcome-guided discovery axis. A two-view model excluding survival served as the unsupervised sensitivity analysis (Fig. 2a; Supplementary Note 3). Supplementary Table S16 reports the factor summary.

Candidate nomination integrated absolute Factor 1 loading ranks with ALS–control abundance, long–short survival contrasts and directional consistency within each RNA view, yielding 19 miRNAs and 16 mRNAs. Supplementary Table S17 records the top-50 loading lists and nominated features. The top-30 sets in Figure 2e–g are display subsets. Controls contributed to the disease contrast; MOFA fitting and survival analyses were restricted to the 45 participants with ALS. Figure 2 and Extended Data Figure 5 Source Data provide loadings and contrast-level results; Supplementary Table S1 collects the additional fold-change summary with its specified strata and significance values.

### miRNA–mRNA target evidence and directional states

Candidate relationships were annotated using miRTarBase, DIANA-TarBase and TargetScan.^45,55,56^ Primary support required an 8mer, 7mer-m8 or 7mer-A1 seed match, a qualifying experimental database record, or both. Sixmer-only matches formed a lower evidence tier. Of 304 possible pairs, 119 met the primary rule; Supplementary Table S18 lists 173 pairs with recorded sequence or database evidence, including 54 sixmer-only pairs. The full 304-pair cross-product is supplied in Figure 2 Source Data I and Extended Data Figure 6 Source Data F.

For participant i and eligible pair k, within-feature z scores assigned one of four states: suppression (miRNA high, mRNA low), de-repression (miRNA low, mRNA high), co-active (both high) or co-silenced (both low). Suppression burden was defined by equation (1):

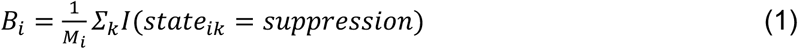

where M_i is the number of eligible pairs for that analysis. Survival-tertile composition and suppression-burden analyses used the 119 prior-supported pairs; complete state-proportion and same-direction control analyses used the full 304-pair cross-product. Group comparisons used two-sided Mann–Whitney tests, continuous survival associations used Pearson or Spearman correlation as specified, and survival curves after median splitting of each participant-level state proportion used two-sided log-rank tests.

### Survival and total-plasma analyses

Kaplan–Meier curves were compared by two-sided log-rank tests, and Cox proportional-hazards models used Efron handling of ties. Three evaluation summaries were kept distinct. For repeated-split displays, 200 stratified 70:30 splits were generated; model weights and dichotomization cut points were estimated in 31-participant training subsets and then applied to all 45 participants with ALS. Curves show the across-split mean with split-to-split variability, and the displayed log-rank P and χ² are the medians of the 200 full-cohort resubstitution tests. Held-out analyses evaluated test participants only. Figure 2m used separate full-cohort univariable Cox models for each standardized continuous molecular or clinical predictor. Row-specific n was determined by complete availability (45 for the three RNA scores, age, sex, C9orf72 status and bulbar onset; 28 for riluzole, wheelchair and bilevel positive airway pressure (BiPAP); 27 for disease duration; 18 for ALSFRS-R decline slope). The slope was multiplied by −1 before standardization so that larger values denote faster decline. Apparent estimates were accompanied by participant-level bootstrap intervals (300 resamples). For split-based analyses, centering, scaling, fitting and cut-point estimation were confined to training data.

For participant i, the unweighted five-miRNA score was defined by equation (2):

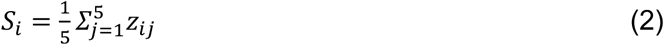

where z_ij is the within-cohort standardized abundance of miRNA j unless a panel specified fitted weights. Empirical survival nulls permuted survival time and censoring status together. For the candidate-reranking null, univariable Cox ranking within the nominated set and score construction were repeated after permutation (Supplementary Note 4). Score definitions, endpoint coding and resampling notation are summarized in Supplementary Note 1.5. Empirical P values used the +1 correction in equation (3):

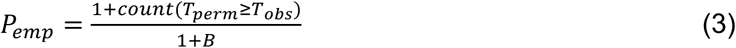

where T_obs is the observed statistic, T_perm the permuted statistic and B the number of permutations. The tail followed the displayed statistic. Horizon-specific AUROC was evaluated at 12, 24, 36 and 54 months. For binary horizon classification, death by the horizon was an event, survival beyond the horizon was a non-event and censoring before the horizon excluded the participant from that endpoint. Full-cohort ROC summaries and cross-validated incremental estimates are reported separately in Figure 3j,k and Supplementary Table S5.

The Magen et al.^31^ total-plasma small-RNA dataset (GSE168714) contained 248 participants with ALS for survival analyses (231 deaths), 243 with NfL measurements and 103 controls for the disease contrast. Counts were renormalized within the present analysis; no published model weights or cut points were reused. All 19 sEV-nominated miRNAs were measurable in total plasma.

For cross-compartment ranking, each candidate j was standardized and fitted in a univariable Cox model in each compartment. Across B = 200 nonparametric bootstrap resamples, the median absolute Cox coefficient was scaled within each compartment to the largest candidate effect, yielding E_EV,j and E_plasma,j. Direction concordance C_j was the fraction of GLAST-positive bootstrap coefficients with the same sign as the full-cohort plasma coefficient. The composite ranking score was defined by equation (4):

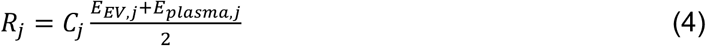

This procedure selected miR-31-5p, miR-4492, miR-93-3p, miR-421 and miR-324-5p (Fig. 3b,c; Supplementary Table S3). Total-plasma survival effects therefore entered refinement of the final five-miRNA panel after the 19-candidate sEV nomination was fixed.

Batch sensitivity was evaluated in the Magen cohort by comparing TMM/log2CPM expression with empirical-Bayes ComBat-adjusted expression.^57^ The recorded batch-adjusted survival analyses included processing batch as a covariate; each model’s covariates and complete-case sample size are specified in the statistical register and Source Data. The five-miRNA score was evaluated with NfL, miR-181 and clinical predictors. Primary survival models used complete cases; classifier-specific imputation is described below. Supplementary Notes 2, 9 and 10 report processing sensitivity, measurement properties and biomarker context.

### Bulk cortical transcriptomic analysis

The 16 candidate mRNAs were evaluated in the New York Genome Center (NYGC) ALS Consortium cortical RNA sequencing (RNA-seq) resource analyzed by Eshima et al.^24^ (GEO GSE153960), comprising 586 profiles from 308 post-mortem donors across ALS, frontotemporal lobar degeneration (FTLD) and control groups and frontal and motor cortical regions (Fig. 4a,b; Extended Data Fig. 9a). Differential-expression contrasts were specified by diagnosis, region and molecular subtype. Contrasts lacking sufficient observations were recorded as not tested/not estimable (NT). Supplementary Table S14 enumerates design matrices, covariates, contrasts and multiple-testing families; Supplementary Data 2 provides complete differential-expression results.

### Single-nucleus cortical analysis

Single-nucleus data were obtained from the Pineda et al.^25^ ALS/FTLD resource (NCBI PRJNA1073234; Synapse syn51105515), comprising 625,973 nuclei from 73 donors across primary motor (Brodmann area 4; BA4) and prefrontal (Brodmann area 9; BA9) cortex. Published donor, diagnosis, region and broad-cell annotations were retained without reclustering. After requiring complete values for variables used here, 527,261 nuclei from 69 donors contributed to Fig. 4 and Extended Data Figs. 9–10. Donor was the biological replicate and inferential unit; nuclei were aggregated to donor-pseudobulk values for inference, consistent with current guidance for multi-sample single-cell studies.^75–77^

Twenty-five prespecified state programs were scored within astrocyte, microglia, oligodendrocyte precursor cell (OPC), oligodendrocyte and vascular parent lineages. Marker sets and parent gates are listed in Supplementary Table S19. Programs were designated a priori as lineage-identity anchors or shared responses; shared interferon, stress, cycling, reactive and inflammatory programs were interpreted only within their prespecified parent lineage. Gate specificity was tested by comparing donor-paired program scores inside versus outside the gate, with 95% intervals from 2,000 bootstrap resamples, two-sided paired Wilcoxon signed-rank tests and BH adjustment across 25 programs (Extended Data Fig. 10a). Cargo–program associations used donor-pseudobulk values and Spearman correlation; panel-specific testing families are stated in Source Data (Fig. 4g,h; Extended Data Fig. 10b). State-program gene sets excluded the five selected cargo genes. Supplementary Table S20 summarizes FDR-significant cargo–program associations; complete values are provided in Source Data Figure 4, sheet G.

### Tissue-bridge and pathway analyses

The 16 mRNA candidates were summarized by tissue-bridge concordance and combined sEV survival support. The primary core required a candidate to exceed the median across all 16 candidates on both axes (Fig. 4c). NPM1, MTMR3, ZBTB16, LAMP1 and CBL met this conjunction. SLC22A8 and STK4 were single-axis boundary cases. Supplementary Table S7 provides the candidate-level component summaries and corrected primary-core labels, while Figure 4 Source Data C retains plotted coordinates. Supplementary Data 1 is assigned to the complete tissue-bridge component matrix.

A separate four-component sensitivity score combined normalized sEV survival evidence, tissue-bridge evidence, bulk classification and gene-constraint/variant context (Fig. 4d). The displayed score ranked NPM1, MTMR3, ZBTB16, LAMP1 and CBL first through fifth, respectively; STK4 ranked sixth and SLC22A8 fourteenth. Figure 4 Source Data D contains the numeric components and ranks. The primary core used the two-axis conjunction; the composite supplied contextual sensitivity analysis. Its annotation resources included gnomAD, ClinVar, the National Human Genome Research Institute–European Bioinformatics Institute (NHGRI–EBI) genome-wide association study (GWAS) Catalog, ALS GWAS resources, Open Targets and Project MinE (Supplementary Table S8). General constraint, variant counts and pathway context were distinguished from direct candidate-level ALS genetic evidence.

Pathway over-representation used Reactome, Gene Ontology and MSigDB Hallmark gene sets^59–61^ with one-sided hypergeometric tests and BH correction within each enrichment family. Supplementary Table S21 records summarized terms and memberships, and Figure 5 Source Data provides the displayed enrichment and module records. Supplementary Data 3 is assigned to predicted miRNA target footprints. Figure 5b shows an FDR-significant subset; Figure 5c summarizes module coverage for cargo mRNAs and predicted miRNA targets, with pathway counts and best enrichment values reported separately. Lineage sectors represent normalized enrichment contributions. The Figure 5a evidence-synthesis network was assembled with EVd3x.^58^

### Machine-learning evaluation

Supporting plasma analyses examined penalized selection and panel-size sensitivity separately from the five-miRNA cross-compartment ranking. Features required log2CPM >1 in at least 50% of samples and more than five unique values. Elastic-net Cox models were evaluated over α ∈ {0.1, 0.3, 0.5, 0.7, 1.0} in 100 bootstrap resamples; stability thresholds were ≥60% selection frequency and ≥80% coefficient-sign consistency. Panels of K = 2–6 were evaluated as independently optimized sensitivity configurations. Supplementary Tables S22 and S23 report these configurations and parameters.

At 12-, 24-, 36– and 54-month horizons, seven classifier families were compared: L2-penalized logistic regression, random forest, gradient boosting, extremely randomized trees, AdaBoost, radial-basis support vector machine and linear discriminant analysis. Models used repeated stratified five-fold cross-validation with 10 repeats (seed 42). Median imputation for missing classifier inputs, together with filtering, scaling, selection and fitting, was confined to training folds; primary survival models used complete cases without imputation. AUROC, balanced accuracy, precision, recall, the F1 score (the harmonic mean of precision and recall), precision–recall and calibration were calculated from out-of-fold predictions, and repeated predictions were aggregated to one value per participant before paired model comparisons. Curve uncertainty in Extended Data Fig. 8 used 400 bootstrap resamples of pooled out-of-fold predictions; nested AUROCs were compared by paired two-sided DeLong tests on participant-aggregated predictions. Supplementary Table S5 reports horizon-dependent incremental performance.

### Interactive reproducibility application

An R Shiny application, the ALS Paired-RNA Explorer (release 1.0.0-paper), was developed as a reproducibility companion. Paper-reproduction mode loads fourteen frozen result tables and reproduces the displayed outputs from fixed results. A separate cohort-scoring workflow accepts de-identified comma-separated value (CSV) files containing the fixed five-miRNA feature set and computes a research-use transfer score without changing feature identities; an optional exploratory paired-RNA workflow uses outcome-guided singular-value decomposition as a lightweight companion analysis. User uploads are processed session-locally. A verification layer recomputes 22 numerical claims from released tables and emits a machine-readable pass/fail manifest. The repository and live application’s uniform resource locator (URL), together with the citable archive’s digital object identifier (DOI), will be provided at journal submission *([AUTHOR TO SUPPLY AT JOURNAL SUBMISSION: application repository URL, Zenodo archive DOI and live deployment URL]*).

### Statistics and reproducibility

All statistical tests were two-sided unless a one-sided enrichment test is specified. Exact P values are reported where available; uncorrected P < 0.05 is described as nominal. Multiple testing was controlled by the Benjamini–Hochberg procedure^78^ within the analysis family stated in the text, legend or Source Data. Exact sample sizes, biological-replicate definitions, summary statistics and tests are reported with the corresponding figures and Source Data. No statistical method was used to predetermine sample size. Missing values were not imputed unless explicitly stated, and complete-case subsets are identified with their n. Donors were the biological replicates for single-nucleus analyses. Random seeds, software versions and model parameters are recorded in the analysis package and Supplementary Table S23.

### Use of generative artificial intelligence and AI-assisted technologies

During manuscript preparation, the authors used the generative artificial intelligence (AI) systems Claude Opus 5 (Anthropic) and ChatGPT (GPT-5.6 Sol; OpenAI) to assist with grammar, sentence clarity, organization and language editing of author-written text. OpenAI Codex was used for code organization, code review and debugging, documentation and repository structure. No generative-AI tool was used as an independent source of scientific evidence or as an independent basis for data interpretation. All analyses, code, figures, interpretations and manuscript text were reviewed and approved by the authors, who take full responsibility for the work.

## Reporting summary

The reporting information for the experimental and analytical procedures is provided in the Methods, figure legends, Source Data and supplementary reporting registers (Supplementary Tables S10 and S11).

## Data availability

### Data generated in this study

Raw sequence data for the 60 discovery participants (one small-RNA and one paired-end transcriptome library per participant) will be deposited in the National Center for Biotechnology Information (NCBI) Sequence Read Archive (SRA); the BioProject accession will be released at journal submission *([AUTHOR TO SUPPLY AT JOURNAL SUBMISSION: SRA BioProject accession]*). The analysis archive includes feature-by-participant raw-count matrices, TMM-normalized log2-CPM matrices for the same 60 participants, a FASTQ-linked de-identified metadata table and dictionary, the fitted 45-ALS MOFA+ object and Supplementary Data 1–4. Exact dates and internal identifiers are retained in a separate controlled-access metadata file governed by institutional review board (IRB), consent and disclosure review. Processed matrices will also be deposited in NCBI Gene Expression Omnibus; the accession will be released at journal submission *([AUTHOR TO SUPPLY AT JOURNAL SUBMISSION: GEO series accession]*). Source Data workbooks contain only observations underlying displayed panels; Supplementary Tables S1–S26 are provided once in a single workbook. Supplementary Table S24 provides the complete data-and-code manifest.

### Data obtained from other studies

The total-plasma small-RNA cohort is from Magen et al.^31^ and is available at GEO under accession GSE168714. Bulk cortical transcriptomes are from Eshima et al.^24^ and are available at GEO under accession GSE153960. Single-nucleus cortical data are from Pineda et al.^25^ and are available at NCBI BioProject PRJNA1073234 with processed matrices at Synapse (syn51105515). These datasets are cited to their original accessions and are not redistributed here; derived tables generated in the present analysis are provided in the Supplementary Information. Reference resources were miRTarBase^56^, DIANA-TarBase^55^, TargetScan^45^, ATtRACT^51^, Reactome, Gene Ontology, MSigDB Hallmark and GENCODE (hg38).

## Code availability

All analysis code will be released at journal submission and archived at Zenodo under a digital object identifier (DOI) *([AUTHOR TO SUPPLY AT JOURNAL SUBMISSION: code repository URL and Zenodo DOI]*). The archive contains panel-level R and Python scripts, the pipeline configuration file specifying every threshold and seed (reproduced as Supplementary Table S23), manuscript build scripts, per-panel sessionInfo() records and an renv lockfile pinning package versions. Analyses were performed in R (edgeR, limma, DESeq2, sva, survival, survminer, MOFA2) and Python (scikit-learn, scipy, statsmodels, pandas, numpy); exact versions are recorded in the lockfile and session records. The same release contains the ALS Paired-RNA Explorer 1.0.0-paper source, its fourteen frozen input tables, the twenty-two-claim verification layer and the session-local cohort-scoring and exploratory modules; the live application URL will be provided at journal submission *([AUTHOR TO SUPPLY AT JOURNAL SUBMISSION: live application URL]*).

## Acknowledgements

We thank the persons living with ALS who enrolled in the National ALS Registry and consented to contribute biospecimens to the National ALS Biorepository, without whom this work would not be possible. Specimens and linked coded data were provided by the National ALS Biorepository, administered by the Agency for Toxic Substances and Disease Registry as a component of the Centers for Disease Control and Prevention National ALS Registry.

The findings and conclusions in this report are those of the authors and do not necessarily represent the official position of the Centers for Disease Control and Prevention or the Agency for Toxic Substances and Disease Registry.

This work was supported by the National Institute of Neurological Disorders and Stroke (R21NS147248). We also thank the control participants and the staff who recruited them. We gratefully acknowledge the Yale Center for Genome Analysis (YCGA) for library preparation and sequencing, the Yale Flow Cytometry Core Facility for nano-flow cytometry, the Yale Keck Biotechnology Resource Laboratory for biophysical and proteomic support, and the Electron Microscopy Facility at the Center for Cellular and Molecular Imaging (CCMI), Yale Research, for negative-stain transmission electron microscopy.

## Author contributions

J.S.W. and D.P. conceived and designed the study. J.S.W. performed the experimental work, developed and carried out the computational analyses, interpreted the results, prepared the figures and tables, and wrote the manuscript. D.P. contributed to study conception and design, supervised the work, obtained funding, and contributed to result interpretation, drafting of the manuscript and its substantial revision. F.B. contributed to experimental data acquisition. U.S. contributed to bioinformatics analysis. All authors reviewed and approved the submitted manuscript. Collectively, the authors assume responsibility for the completeness and accuracy of the data.

## Competing interests

J.S.W. and D.P. are co-founders of Eatsrock Dx Inc. and hold equity in the company. F.B. and U.S. declare no competing interests.

---

**Extended Data Fig. 1.**
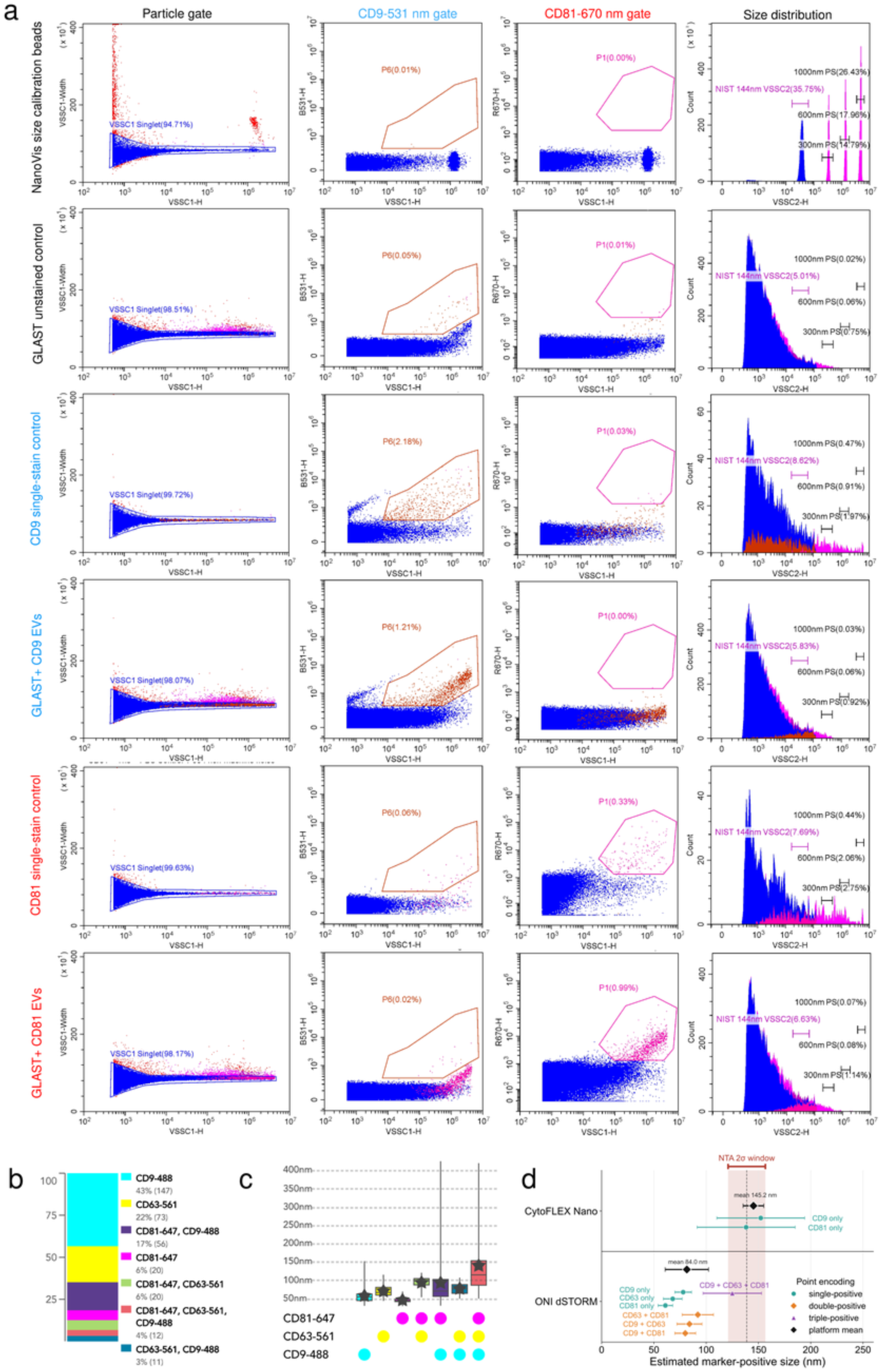
| Orthogonal single-particle measurements confirm tetraspanin heterogeneity of GLAST-positive sEVs. **a**, Representative CytoFLEX Nano gating sequence. Rows show NanoVis size-calibration beads, GLAST unstained control, CD9 single-stain control, GLAST-positive CD9-labelled sEVs, CD81 single-stain control and GLAST-positive CD81-labelled sEVs. Columns show the violet side-scatter singlet-particle gate, CD9-531-nm gate (P6), CD81-670-nm gate (P1) and side-scatter size distribution referenced to 144-nm NIST and 300-, 600– and 1,000-nm polystyrene standards. Every percentage is calculated relative to the parent gate shown in that row and column; unstained and single-stain controls define fluorescence gates. **b,** Tetraspanin phenotypes resolved by three-colour dSTORM (CD9-488, CD63-561 and CD81-647) among 339 localized particles: CD9 alone, 43% (147); CD63 alone, 22% (73); CD81 + CD9, 17% (56); CD81 alone, 6% (20); CD81 + CD63, 6% (20); triple positive, 4% (12); and CD63 + CD9, 3% (11). **c,** dSTORM-estimated particle size by tetraspanin phenotype. Boxes show the median and interquartile range with individual particles overlaid; the marker key identifies single-, double– and triple-positive phenotypes. **d,** Cross-platform comparison of marker-positive size estimates. Point shape denotes single-, double-or triple-positive phenotype and diamonds denote platform means (CytoFLEX Nano, 145.2 nm; ONI dSTORM, 84.0 nm). CytoFLEX Nano values are side-scatter optical size estimates, whereas ONI dSTORM values are localization-defined marker-positive size estimates. The shaded band is the 2-s.d. interval from nanoparticle tracking and is shown as the hydrodynamic-size reference.

**Extended Data Fig. 2.**
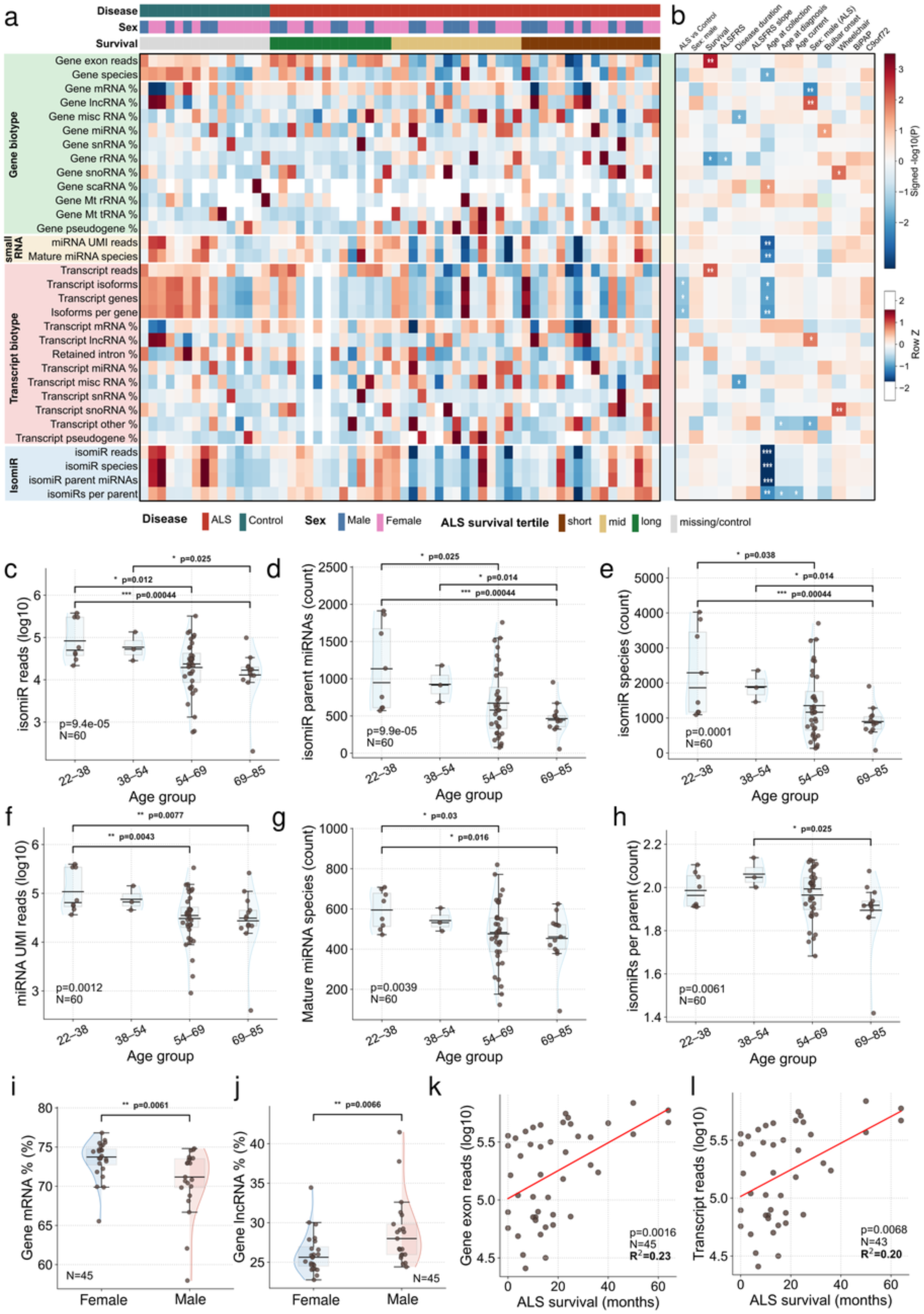
| Demographic and technical axes dominate global GLAST-positive sEV RNA variation. **a**, Sample-level heatmap of 32 gene-, mature-miRNA-, transcript– and isomiR-level cargo measures across 60 GLAST-positive sEV-RNA profiles (45 ALS and 15 controls). Values are scaled within each measure; top annotations show diagnosis, sex and ALS survival tertile. **b,** Association atlas relating the 32 composition measures to ALS versus control status, sex, survival, ALSFRS-R, disease duration, ALSFRS-R slope, age at collection, age at diagnosis, current age among ALS participants, sex among ALS participants, bulbar onset, wheelchair use, BiPAP and C9orf72 status. Continuous variables were tested by two-sided Spearman correlation, binary variables by two-sided Mann–Whitney tests and categorical variables by Kruskal–Wallis tests. Binary signs are ALS minus control, female minus male, yes minus no or present minus absent. Symbols denote ∼P < 0.10, *P < 0.05, **P < 0.01 and ***P < 0.001; BH-adjusted results for the complete family are provided in Source Data. **c–h,** Cargo-complexity measures across age-at-collection groups (22–38, 38–54, 54–69 and 69–85 years): total isomiR reads (c), detected isomiR parent miRNAs (d), detected isomiR species (e), mature-miRNA UMI reads (f), detected mature-miRNA species (g) and isomiRs per parent miRNA (h). Points are participants; distributions and summary boxes are shown for each age group. The P value at lower left is the two-sided Kruskal–Wallis test; brackets show two-sided Dunn pairwise tests with Holm adjustment across the displayed comparisons. **i,j,** Gene-level mRNA (i) and lncRNA (j) read fractions by sex among participants with ALS (n = 45); two-sided Mann–Whitney P = 0.00612 and 0.00655, respectively. **k,l,** Gene-exon read depth (k; Spearman rho = 0.456, P = 0.00165, n = 45) and transcript read depth (l; rho = 0.407, P = 0.00683, n = 43) versus ALS survival. Red lines and displayed R² values are ordinary-least-squares visual summaries; inference uses the stated two-sided Spearman tests.

**Extended Data Fig. 3.**
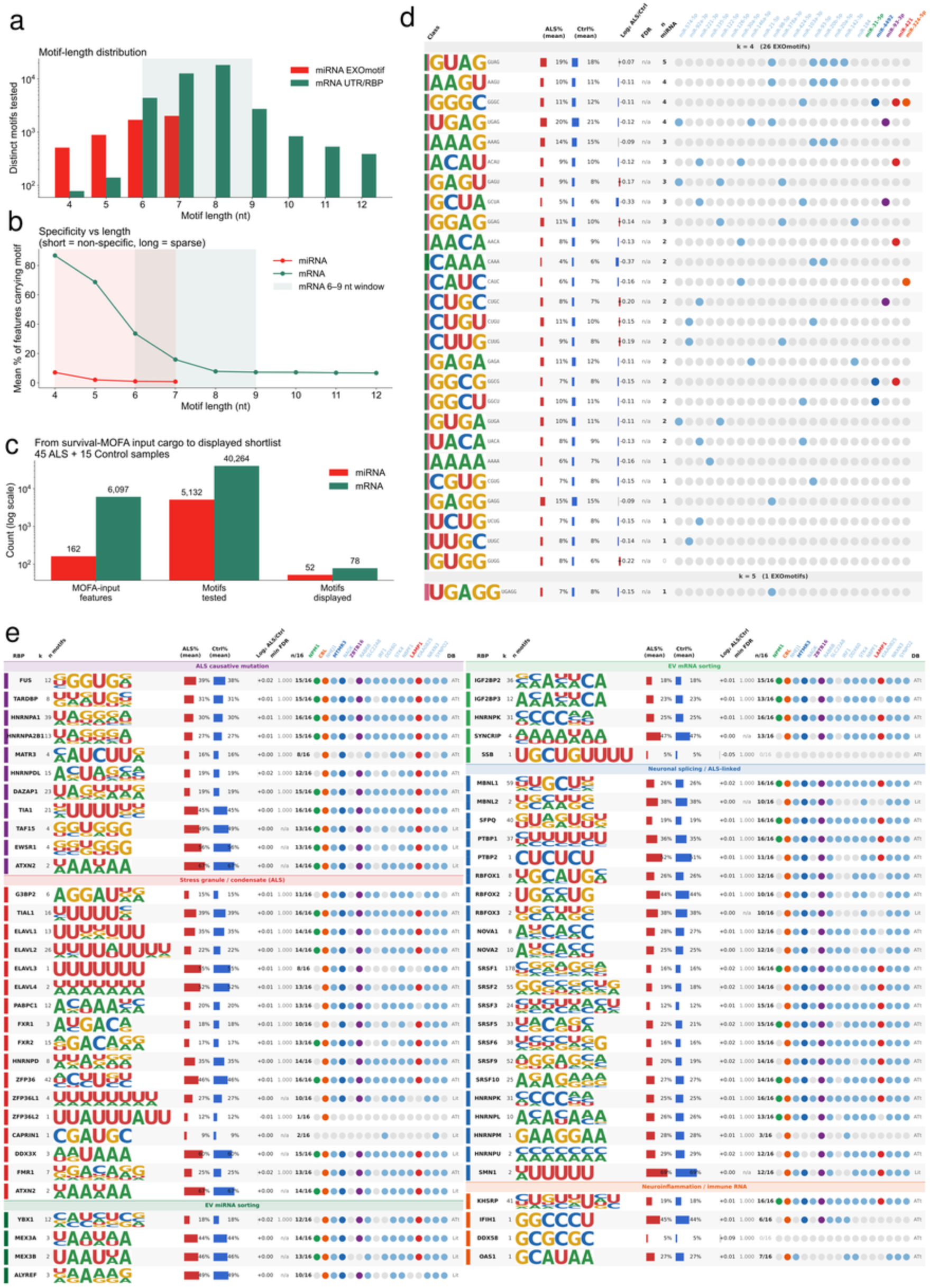
| GLAST-positive sEV RNA retains RBP-associated sequence motifs across miRNA and mRNA cargo. **a**, Number of distinct motifs evaluated at each sequence length in the miRNA EXOmotif and mRNA 3′-UTR/RNA-binding-protein (RBP) universes; the y axis is logarithmic. In total, 5,132 miRNA motifs were evaluated across the motif-analysis universe of 162 eligible miRNAs and 40,264 mRNA motifs across 6,097 protein-coding genes with representative 3′ UTRs. **b,** Mean maximum motif prevalence in ALS or control GLAST-positive sEV-RNA profiles by motif length. Shading marks the prespecified 4–7-nt miRNA EXOmotif and 6–9-nt mRNA UTR/RBP windows. **c,** Feature flow to the displayed motif atlas: 162 miRNAs and 6,097 mRNAs defined the motif-analysis universe; these counts are distinct from the 187 miRNAs and 374 mRNAs entering the outcome-informed MOFA+ model. The displayed atlas retained 52 miRNA entries (27 unique sequences) and 78 mRNA entries. **d,** miRNA EXOmotif atlas. Rows are motif logos; columns report motif class, ALS and control prevalence, log2 ALS/control prevalence ratio, BH FDR, number of the 19 candidate miRNAs containing each motif and candidate-specific containment. **e,** ATtRACT– and literature-supported RBP motifs in 3′ UTRs of the 16 candidate mRNAs, grouped by ALS-causative mutation, stress-granule/condensate, EV-miRNA sorting, neuronal-splicing/ALS-linked or neuroinflammation/immune-RNA categories. Columns show RBP, motif length/count, logo, prevalence, log2 ratio, FDR and candidate containment; dots indicate motif-sequence containment. ALS–control prevalence used two-sided Fisher exact tests with Benjamini–Hochberg correction within motif families; all computable displayed tests had FDR = 1, with nominal P values ≥ 0.05. The panel provides RBP-associated sequence context for the recovered cargo; n/a denotes non-estimable tests.

**Extended Data Fig. 4.**
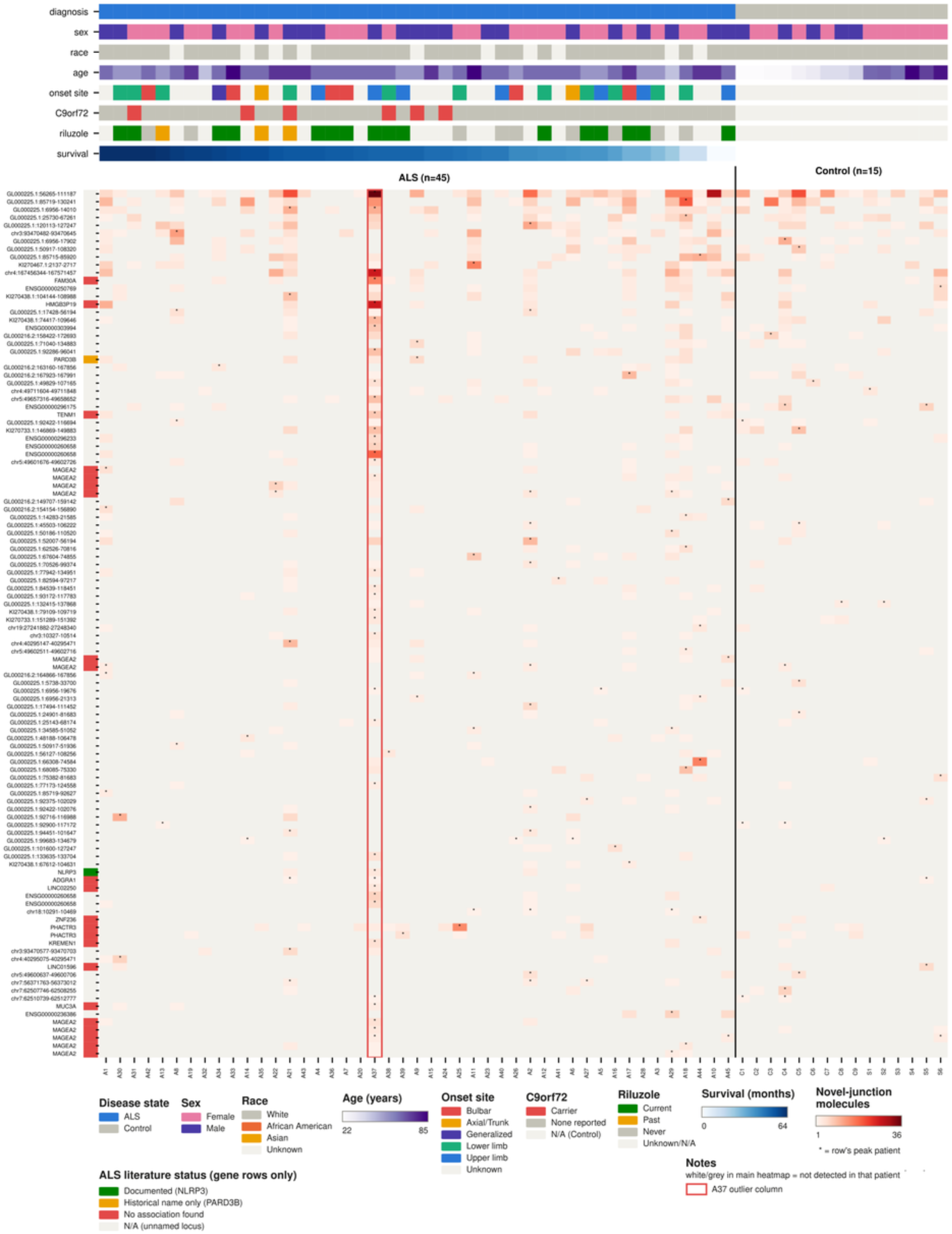
| Low-input GLAST-positive sEV RNA preserves recurrent transcript junction structure. Heatmap shows molecule counts for 110 recurrent unannotated junctions with canonical splice-site motifs across all 60 GLAST-positive sEV-RNA profiles (45 ALS and 15 controls). ALS samples are ordered by survival and controls shown separately. Top tracks denote diagnosis, sex, race, age, onset site, C9orf72 status, riluzole use and survival. Colour gives junction-supporting molecule count; white or grey indicates no detection. An asterisk marks the participant with the maximum count for that junction and serves as a location marker. The red outline identifies participant A37, whose junction-spanning molecule complexity was approximately twice that of the next-highest ALS sample despite mid-range sequencing depth and mapping rate. The left annotation summarizes the scoped literature status of named loci; NLRP3 had ALS-related precedent, whereas PARD3B is retained as a historical-name caution. Sixty-one of 110 recurrent junctions mapped to unplaced or alternative GRCh38 scaffolds. Published KCNQ2-, STMN2– and UNC13A-associated TDP-43 cryptic junctions were queried separately using the stringent molecule filters described in Methods; the depth-limited audit yielded zero qualifying molecules at those reported coordinates. The panel therefore reports the junction inventory supported at the obtained sequencing depth.

**Extended Data Fig. 5.**
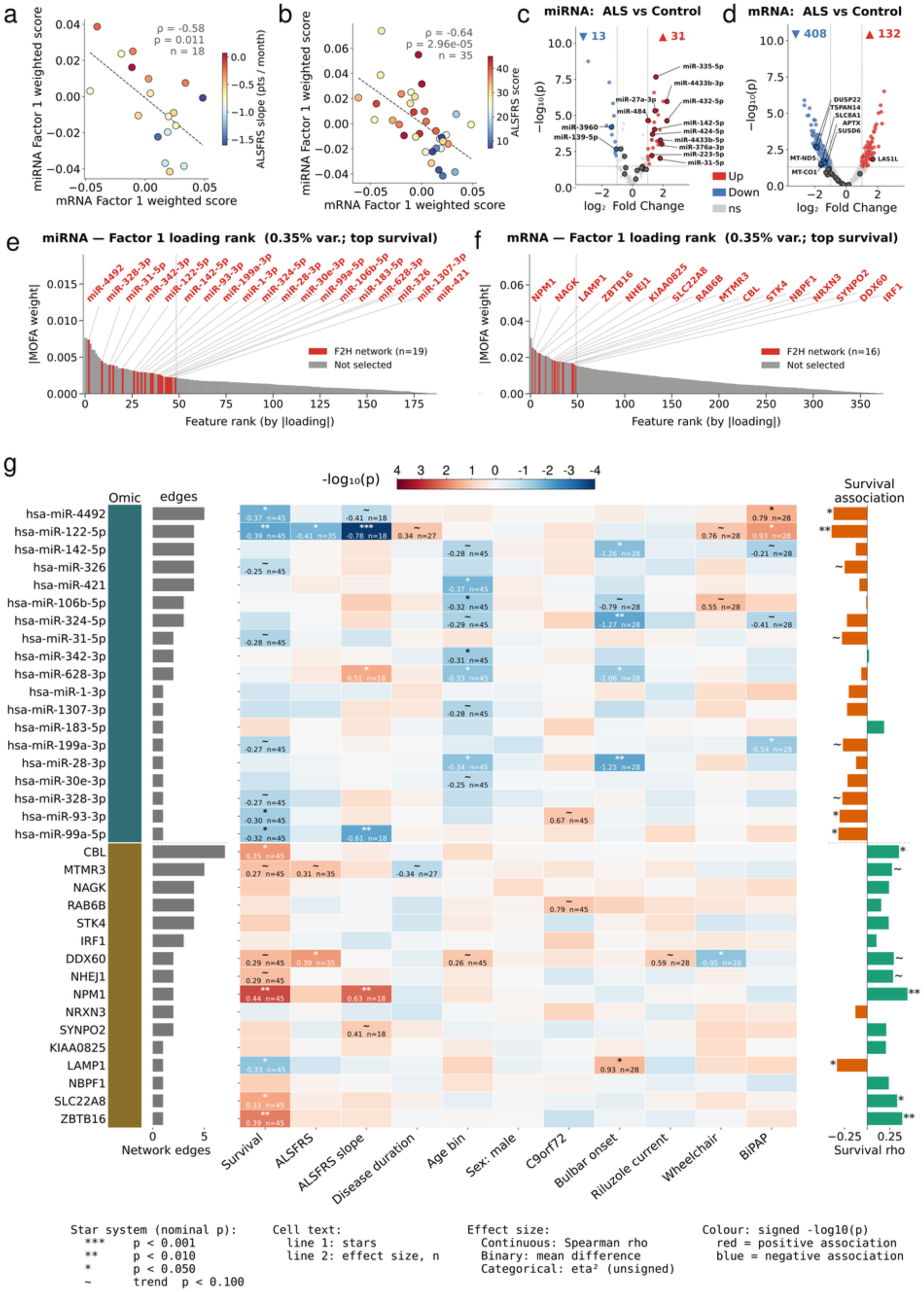
| Clinical overlays and feature-level associations characterize the outcome-guided Factor 1 program. **a,b**, Factor 1 miRNA score versus mRNA score in participants with recorded ALSFRS-R slope (a; Spearman rho = –0.58, two-sided P = 0.011, n = 18) or ALSFRS-R score (b; rho = – 0.64, P = 2.96 × 10^-5, n = 35). Point colour denotes the corresponding clinical measure; neither ALSFRS-R endpoint was used to fit the survival view. **c,d,** Differential abundance of miRNAs (c) and mRNAs (d) in ALS versus control GLAST-positive sEV-RNA profiles. Dashed lines mark nominal P = 0.05 and |log2 fold change| = 1 display thresholds. Thirteen miRNAs were lower and 31 higher in ALS; 408 mRNAs were lower and 132 higher. Labelled points are prioritized features. **e,f,** Absolute Factor 1 loading rank across all 187 miRNAs (e) and 374 mRNAs (f) entering the outcome-informed MOFA+ model. The 19 miRNA and 16 mRNA candidates are highlighted, and the vertical dashed line marks the top-50 loading prefilter. These are the outcome-guided loading ranks used for candidate nomination. **g,** Association matrix relating the 19 candidate miRNAs and 16 candidate mRNAs to survival, ALSFRS-R score, ALSFRS-R slope, disease duration, age band, sex, C9orf72 status, bulbar onset, current riluzole use, wheelchair use and BiPAP. Continuous variables use two-sided Spearman tests, binary variables two-sided Mann-Whitney tests and categorical variables Kruskal-Wallis tests. Heatmap colour is signed –log10 P; cell text gives nominal significance, effect size and available n. Effect sizes are Spearman rho for continuous variables, mean difference for binary variables and eta-squared for categorical variables. Symbols denote ∼P < 0.10, *P < 0.05, **P < 0.01 and ***P < 0.001. The left bars show network degree and the right bars survival rho.

**Extended Data Fig. 6.**
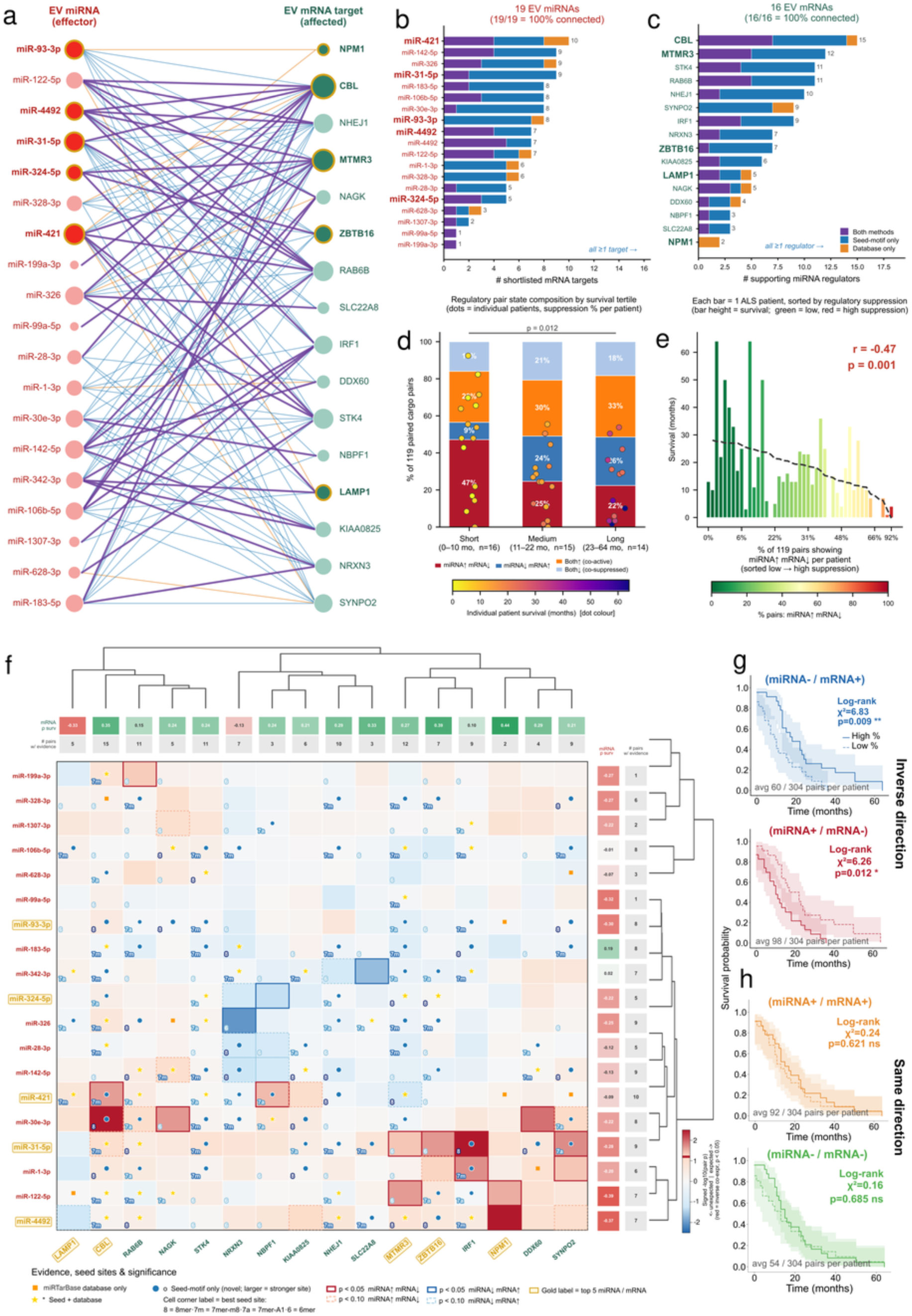
| Reciprocal miRNA–mRNA states are associated with survival. **a**, Bipartite network of the 119 prior-supported relationships among 19 candidate miRNAs and 16 candidate mRNAs; all 35 features had at least one supported partner. Node size reflects degree, gold outlines mark the five highest-priority features in each RNA class and edge colour denotes seed-only, database-only or combined support. **b,c,** Number of supported mRNA partners per miRNA (b; range 1–10) and miRNA regulators per mRNA (c; range 2–15), partitioned by evidence class. **d,** Participant-level composition of four directional states across survival tertiles: short, 0–10 months (n = 16); intermediate, 11–22 months (n = 15); and long, 23–64 months (n = 14). States are suppression (miRNA high/mRNA low), de-repression (miRNA low/mRNA high), co-active (both high) and co-silenced (both low), defined from within-feature z scores across the 119 supported pairs. Suppression averaged approximately 47%, 25% and 22% across tertiles; short versus long, two-sided Mann–Whitney P = 0.012. **e,** Survival after ordering participants by suppression burden (Pearson r = −0.47, P = 0.001, n = 45). **f,** Complete 19 × 16 pair matrix. Fill gives pairwise expression association; symbols denote seed/database evidence and outlines nominal inverse-or de-repression-direction associations. **g,h,** Kaplan–Meier analyses after median dichotomization of participant pair-state proportions across all 304 candidate pairs. Inverse-direction states separated survival (de-repression P = 0.00899; suppression P = 0.0123), and same-direction states yielded P = 0.621 (co-active) and P = 0.685 (co-silenced). Two-sided log-rank tests; n = 45, 45 events.

**Extended Data Fig. 7.**
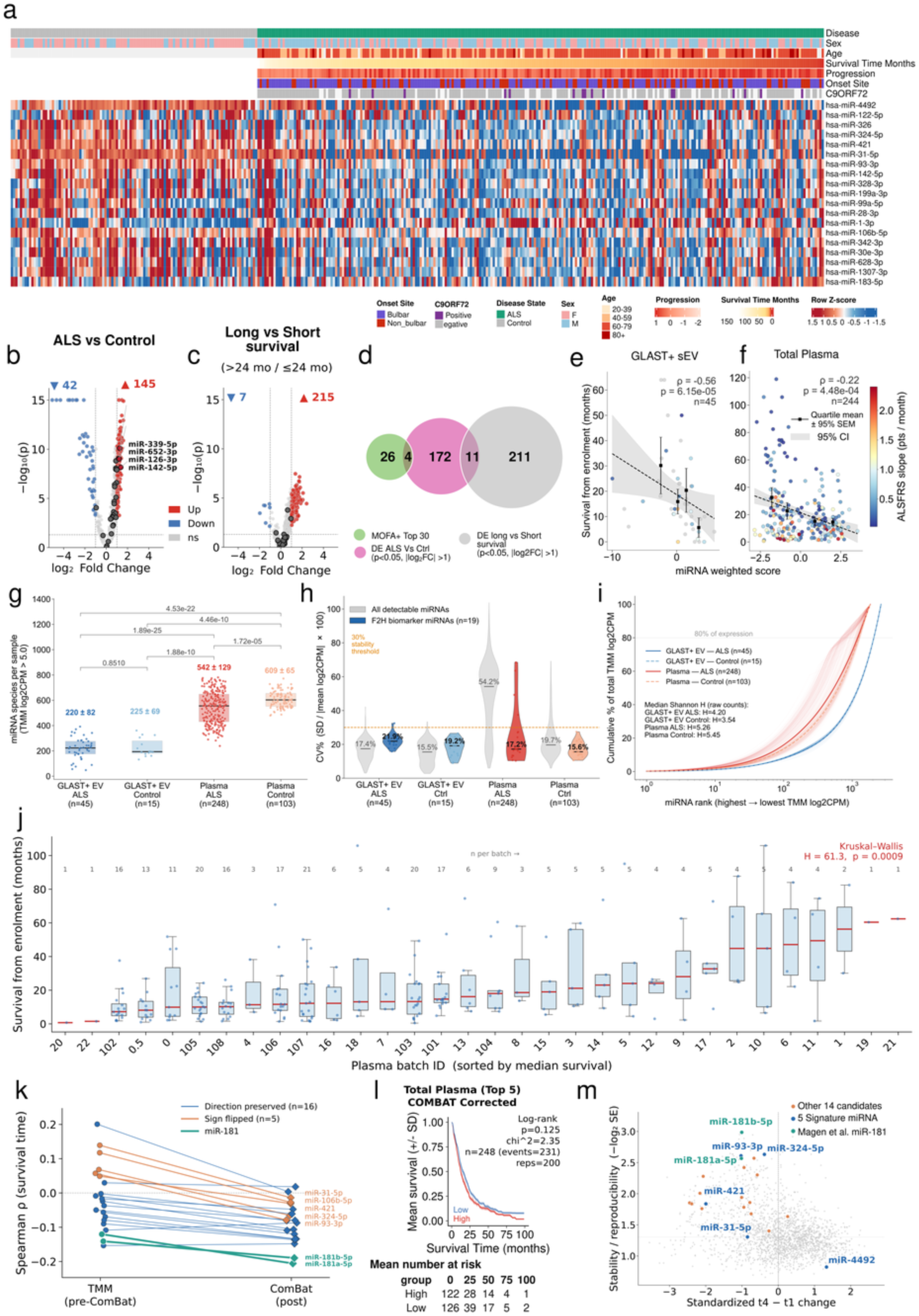
| The transferred miRNA program is measurable in total plasma and shows processing-dependent survival associations. Unless otherwise stated, panels use Magen et al.^31^ total plasma (248 ALS, 103 controls; 231 ALS deaths; 243 ALS NfL measurements). **a,** Row-scaled abundance of the 19 sEV-nominated miRNAs with clinical annotations. **b,c,** Differential abundance for ALS versus controls (b) and long (>24 months) versus short survival (c); dashed lines mark nominal P = 0.05 and |log2 fold change| = 1. **d,** Overlap among MOFA+ top-30, plasma disease and plasma survival display sets. **e,f,** Five-miRNA score versus survival in GLAST-positive sEVs (e; n = 45, Spearman ρ = −0.56, P = 6.15 × 10−5) and plasma (f; n = 244, ρ = −0.22, P = 4.48 × 10−4). **g,** Detectable miRNAs per sample at log2CPM >5 in sEV ALS, sEV controls, plasma ALS and plasma controls; values above groups are mean ± s.d. **h,** Interparticipant coefficient of variation (CV) of log2CPM (100 × s.d./mean) for all detectable miRNAs and the 19 candidates; horizontal line, 30% CV. **i,** Cumulative ranked abundance; dashed line, 80% cumulative expression. Median Shannon diversity was 4.20 in sEV ALS and 5.26 in plasma ALS; control summaries are shown in the panel. **j,** Survival by processing batch; two-sided Kruskal–Wallis H = 61.3, P = 0.0009. **k,** Candidate-wise survival correlations before and after ComBat among 122 participants with complete expression, survival and processing-batch information; 10 directions were preserved and 9 changed sign. **l,** Repeated-split Kaplan–Meier summary for the ComBat-corrected score (n = 248, 231 events, 200 splits; log-rank P = 0.125). **m,** Longitudinal stability summary (−log2 standard error) versus standardized t4-to-t1 change in 22 participants; groups distinguish background miRNAs, remaining sEV candidates, the five-miRNA signature and miR-181a-5p/miR-181b-5p comparators.

**Extended Data Fig. 8.**
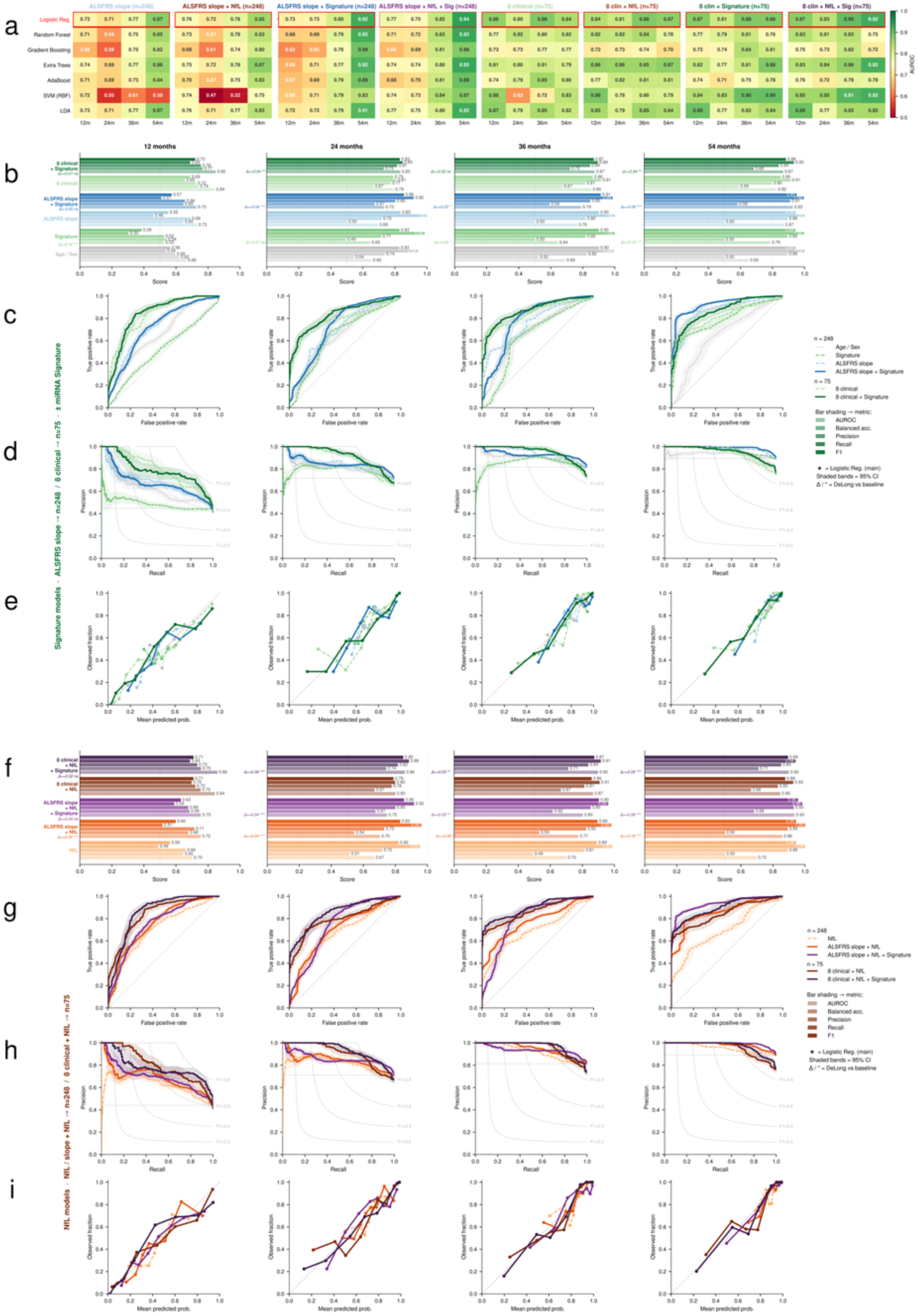
| RNA adds horizon-dependent prognostic information across classifiers and clinical baselines. Analyses use the Magen et al.^31^ total-plasma cohort (GEO GSE168714). **a,** AUROC heatmaps for seven classifiers at 12-, 24-, 36– and 54-month survival horizons. The first four model blocks use ALSFRS-R slope, NfL, the five-miRNA signature and their combinations in the molecular cohort; the final four use an eight-variable clinical model and additions of NfL and/or the signature in the complete-clinical subset (n = 75). **b,** Signature-focused summary of AUROC, balanced accuracy, precision, recall and F1. Bars summarize classifier performance, the star denotes the prespecified logistic-regression result and Δ/P annotations compare nested models by paired two-sided DeLong test on participant-aggregated out-of-fold predictions. **c–e,** Corresponding ROC curves with 95% bootstrap bands (c), precision–recall curves with bootstrap bands and F1 iso-curves (d), and calibration curves (e). **f–i,** Equivalent summaries for NfL-focused comparisons. At each horizon, death by the horizon is an event, survival beyond the horizon a non-event and censoring before the horizon excludes that participant. Models used median imputation where required, training-fold standardization and repeated stratified five-fold cross-validation with 10 repeats (seed 42); curve uncertainty used 400 bootstrap resamples of pooled out-of-fold predictions.

**Extended Data Fig. 9.**
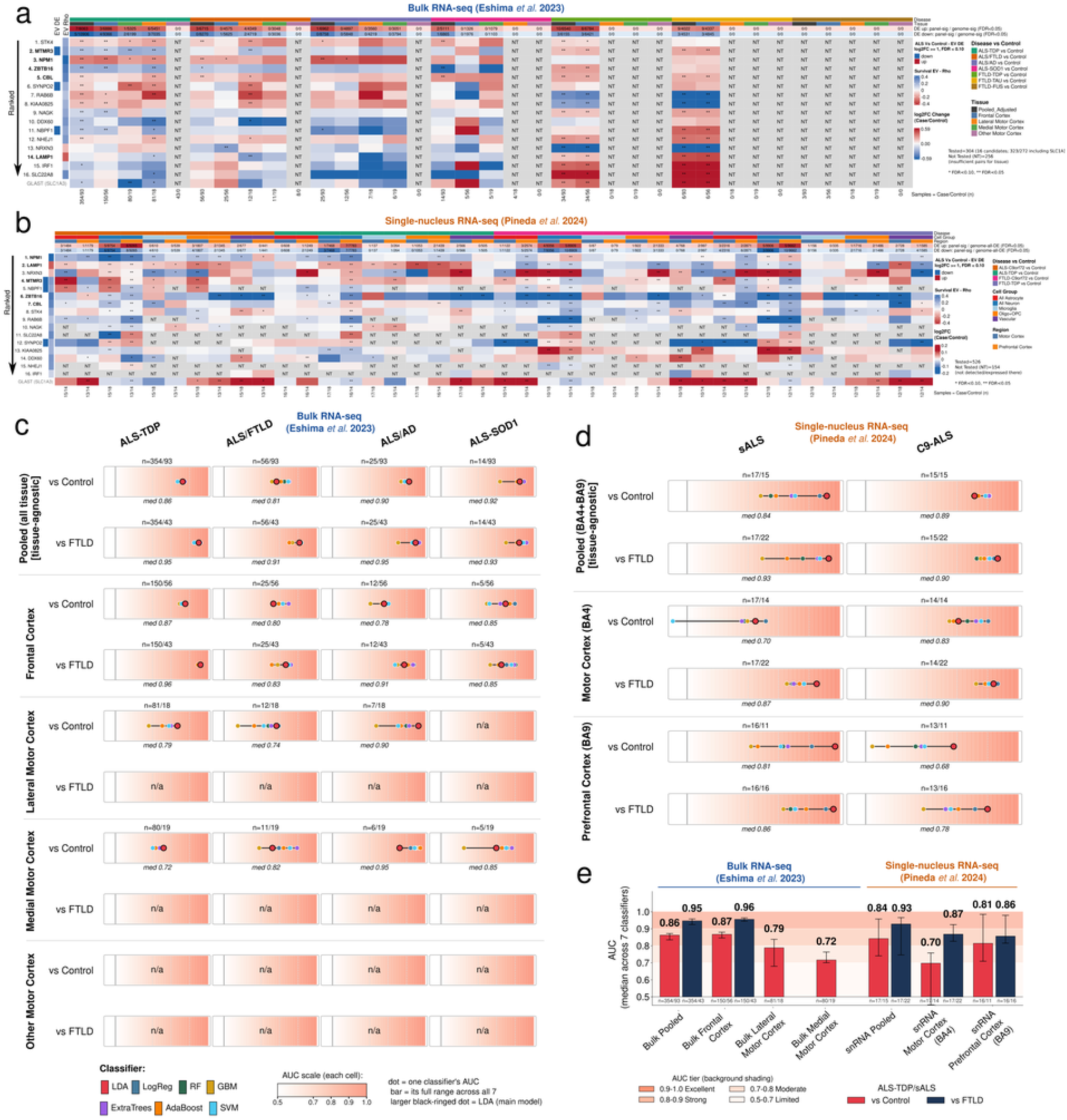
| External cortical datasets contextualize the blood-nominated mRNA program across disease and region. **a**, External bulk RNA-sequencing differential-expression audit for 16 candidate mRNAs plus SLC1A3 across 35 disease-by-region contrasts in the NYGC ALS Consortium resource analyzed by Eshima et al.^24^ (586 profiles from 308 donors). Rows are ordered by sEV evidence; fill denotes log2 fold change; *FDR < 0.10, **FDR < 0.05; NT, contrast unavailable for estimation. **b,** Corresponding donor-pseudobulk audit in the Pineda et al.^25^ single-nucleus resource across diagnosis, cortical region and broad cell class. Donor is the inferential unit. **c,d,** Classification using the selected five-mRNA signature (NPM1, MTMR3, ZBTB16, LAMP1 and CBL) across bulk-tissue contrasts (c) and single-nucleus donor-pseudobulk contrasts (d). Coloured dots, AUROC for seven classifiers; horizontal lines, classifier range; enlarged black-ringed points, prespecified linear-discriminant-analysis result; case/comparator n is printed in each cell. **e,** Median AUROC across classifiers for bulk pooled and regional comparisons and single-nucleus pooled BA4 + BA9, BA4 and BA9 comparisons against controls or FTLD. Error bars show across-classifier spread; background bands denote AUROC tiers. These comparisons characterize the selected core within the tissue resources used for its refinement. Pooled contexts anchor interpretation; small regional or subtype cells are descriptive.

**Extended Data Fig. 10.**
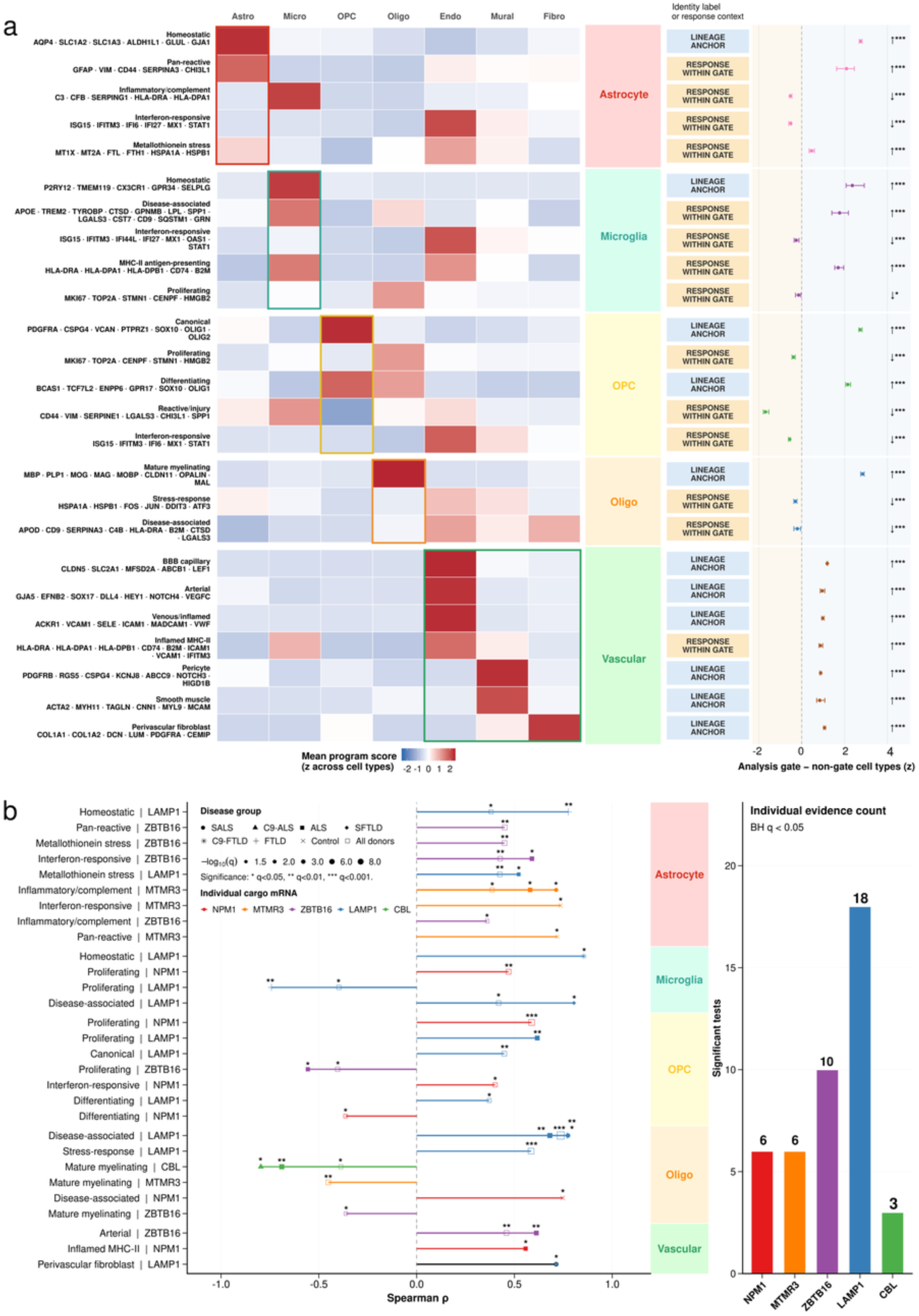
| Prespecified lineage gating localizes the cortical programs coupled to the five-gene cargo score. **a**, Localization of the 25 state programs used in Fig. 4 across seven broad cell classes. Heatmap colour is the mean program score z-scaled across cell classes; coloured boxes mark the prespecified parent gate (astrocyte, microglia, OPC, oligodendrocyte or pooled endothelial + mural + fibroblast vascular cells). Adjacent labels classify each program as a lineage-identity anchor or a response interpreted within its parent gate. At right, points and horizontal bars show the donor-paired difference between analysis-gate and non-gate cell classes with 95% bootstrap intervals (2,000 resamples); P values are from two-sided paired Wilcoxon signed-rank tests and q values are Benjamini–Hochberg adjusted across 25 programs. Shared interferon, inflammatory, reactive, stress and cycling programs are interpreted only within the prespecified gate. **b,** Significant donor-pseudobulk Spearman associations between the five selected cargo mRNAs and 25 gated state programs across displayed disease groups. Each point is one gene–program–disease association; x position, Spearman ρ; point shape, disease group; colour, cargo gene; size, −log10 q. Only associations passing q < 0.05 in the 1,000-test family are displayed. Forty-three associations met this threshold: NPM1, 6; MTMR3, 6; ZBTB16, 10; LAMP1, 18; CBL, 3. State-program gene sets exclude the five selected cargo genes.

