## Supplemental Information for "Paired RNA profiling of circulating small extracellular vesicles links survival in ALS to reactive glial - vascular programs"

### Supplementary Note 1. Analysis definitions and reproducibility

#### 1.1 Reproducibility implementation

Supplementary Tables S14 and S23 record feature filters, penalized-selection settings, stability thresholds, panel-size analyses and classifier parameters. Seven-classifier comparisons use five-fold cross-validation repeated ten times (seed 42), with fold-dependent imputation and scaling estimated in training data. Participant-aggregated predictions support paired model comparisons; uncertainty calculations are identified separately where pooled repeated predictions were used.

#### 1.2 Score definitions, endpoints and resampling notation

The following definitions expand the conventions described in the main Methods. Let $z_{ij}$ denote the standardized abundance of feature j in participant i. For held-out analyses, centering and scaling parameters are estimated in the training subset and applied to the test subset. The unweighted five-miRNA score is the arithmetic mean of the five standardized abundances:

$$S_i = \frac{1}{5}\Sigma_{j=1}^{5} z_{ij}$$

The suppression-state burden is the fraction of eligible pairs with miRNA above and mRNA below their standardized reference values. $M_i$ denotes the number of eligible pairs. The panel-specific universe is either 119 supported pairs or all 304 candidate pairs.

$$B_i = \frac{1}{M_i}\Sigma_k I(state_{ik} = suppression)$$

The empirical permutation P value includes the observed configuration through the +1 correction. B denotes the number of randomizations. Null statistics are counted in the tail appropriate to the observed statistic. Survival time and event status are permuted together; score-level tests retain selected features, whereas the candidate-reranking test repeats univariable Cox ranking within the 19-candidate set.

$$P_{emp} = \frac{1+count(T_{perm} \geq T_{obs})}{1+B}$$

Cross-compartment prioritization combines scaled bootstrap Cox-effect magnitudes in the sEV compartment and total plasma, multiplied by direction concordance $C_j$. This feature-ranking score is distinct from the participant-level expression score above.

$$R_j = C_j \frac{E_{EV,j}+E_{plasma,j}}{2}$$

At horizon τ, recorded death by τ defines an event, survival beyond τ defines a non-event, and censoring before τ excludes the participant from the horizon-eligible binary analysis. Repeated full-cohort Kaplan–Meier displays, held-out survival analyses and participant-aggregated out-of-fold classifier comparisons are separate evaluation summaries. Sample sizes, score configurations and resampling units are specified by panel in the statistical register.

### Supplementary Note 2. Preprocessing, batch structure and depth sensitivity

The analyses used raw counts, TMM-normalized log2CPM, variance-stabilized expression, ComBat-adjusted expression and standardized MOFA+ inputs. Supplementary Table S14 maps each panel to its recorded input and test. The data manifest lists the matrices and fitted objects for release.

Processing batch covaried with survival in total plasma (Kruskal–Wallis H = 61.3, P = 9 × $10^{-4}$; Extended Data Fig. 7j). The 19-candidate ComBat source table and statistical summary record ten preserved and nine changed survival-correlation signs, with a median correlation shift of −0.093. Repeated-split five-miRNA separation after correction gave P = 0.125 (Extended Data

Fig. 7k,l). These results establish a material dependence on preprocessing in a cohort with aligned batch and outcome structure.

Normalized-expression and ComBat results are reported together, with batch adjustment specified for each model. Their divergence motivates randomized processing and prespecified normalization in prospective evaluation (Note 10).

Discovery mRNA sequencing depth correlated with survival (ρ = 0.456, P = 0.0017; Extended Data Fig. 2k). Adjustment for gene-exon read depth preserved associations for the miRNA, mRNA and combined scores (hazard ratios per s.d. 1.82, 3.13 and 2.86; P = 0.0092, $1.10 \times 10^{-4}$ and $6.00 \times 10^{-5}$; n = 45), and transcript-depth adjustment produced similar estimates in 43 complete cases. Depth normalization also attenuated the survival association of unannotated-junction burden (Extended Data Fig. 4).

### Supplementary Note 3. Unsupervised integration sensitivity analysis

An outcome-free two-view MOFA model using the same miRNA and mRNA matrices returned eight factors. F4 had a nominal survival association (ρ = −0.302, P = 0.0439; q = 0.278 after Benjamini–Hochberg correction), whereas F1 accounted for 24.3% of mRNA variance with little survival association. None of the eight factors met the multiplicity-adjusted threshold for survival. These results distinguish the dominant unsupervised axes from the structure prioritized by outcome-guided discovery (Supplementary Table S16).

### Supplementary Note 4. Permutation and pairing controls

Three permutation analyses test fixed-score association, outcome-based candidate ranking and the contribution of matched participant identity. Each uses the null procedure specified below.

#### 4.1 Score-level permutation

Survival time and censoring status were jointly permuted against the locked five-miRNA score, preserving the observed censoring structure.

Across 2,000 iterations, the null median C-index was 0.514 versus 0.570 for the locked score (empirical P = 0.002). The observed log-rank $\chi^2$ of 24.74 exceeded the null median of 0.42 and every permutation (P < 0.0005).

This test conditions on the selected five features. Section 4.2 additionally repeats candidate ranking under permuted outcomes.

#### 4.2 Outcome permutation with candidate reranking

For each of 1,000 iterations, survival time and censoring status were permuted together, the 19 candidates were ranked by univariable Cox |z|, the top five were selected and the score was rebuilt.

| Metric | Value |
|---|---|
| Observed C-index, selection rerun | 0.5740 |
| Null median | **0.5435** |
| Null 95th percentile | 0.5681 |
| Null maximum | 0.5969 |
| Empirical P | **0.026** |

The reranking null had a median C-index of 0.5435, a 95th percentile of 0.5681 and a maximum of 0.5969. The observed C-index under this procedure was 0.5740 (empirical P = 0.026). Its difference from the fixed-score value of 0.570 reflects re-derivation of the five-feature panel.

This null repeats selection by univariable Cox |z| within the nominated set. It leaves the preceding discovery steps fixed and uses a different ranking rule from the composite bootstrap concordance procedure for the reported transfer panel.

#### 4.3 Participant-pair destruction

Participant identity was permuted between the miRNA and mRNA matrices and the cross-layer weighted-score correlation was recalculated 10,000 times.

The observed correlation was 0.7384 (n = 45), compared with a null mean of 0.000, a 95th percentile of |r| of 0.2917 and a maximum of 0.6020; no permutation reached the observed value (empirical P = 1/10,001). This test shows that the fitted scores retain participant-specific cross-layer structure. Factor fitting and candidate selection remain fixed and are not retested by this null.

### Supplementary Note 5. Feature selection and analysis thresholds

The candidate set comprised 19 miRNAs and 16 mRNAs nominated from the outcome-guided discovery model. Subsequent prioritization used the recorded cross-compartment ranking for miRNAs and the conjunction and tissue-bridge criteria for mRNAs. Exact filters, thresholds and component scores are given in Supplementary Tables S3, S4, S7, S8 and S23.

Figure 2c,d uses >18 versus ≤18 months for its discovery survival contrast, as labeled in the source figure. Some total-plasma displays use 24 months. The additional fold-change summary in Supplementary Table S1 uses a distinct, explicitly defined ≥18 versus <18-month contrast in both compartments: 18 versus 27 participants in discovery and 99 versus 145 in plasma. Its plasma strata include 244 participants with available survival time. Continuous survival models retain event/censoring information; each dichotomized display is interpreted according to its own threshold and inclusion rule.

Volcano and overlap displays use nominal P < 0.05 together with |log2 fold change| ≥ 1 for visual classification. Nominal P values and Benjamini–Hochberg-adjusted FDRs are reported separately. Supplementary Table S1 retains the FDR values supplied with its differential-abundance records, alongside the effect-size estimates.

The 19 sEV-nominated miRNAs were ranked by a bootstrap score that combines scaled Cox effects in both compartments with sEV-to-plasma direction concordance. This selected miR-31-5p, miR-4492, miR-93-3p, miR-421 and miR-324-5p.

Bootstrap direction concordance distinguished candidates with similar effect magnitudes. miR-324-5p had 99% concordance, whereas miR-1-3p had zero concordance despite a plasma association (P = 0.00199). The original Magen analysis also excluded miR-1-3p from prognostic testing because of longitudinal variability. Concordance refers to Cox-effect orientation; ALS–control fold-change direction and raw survival-time correlation are separately reported quantities.

The tissue core required each of the 16 mRNA candidates to exceed the candidate median on both the sEV-survival and tissue-bridge axes. NPM1, MTMR3, ZBTB16, LAMP1 and CBL met this conjunction (Fig. 4c; Supplementary Table S7). A separate four-component analysis incorporating gene-constraint/variant context, bulk classification, tissue bridging and survival support ranked the same five genes highest; STK4 ranked sixth and SLC22A8 fourteenth in the displayed score (Fig. 4d; Supplementary Table S8). The conjunction defines the primary core; the composite provides a sensitivity ranking.

Repeated survival displays used 200 stratified 70:30 splits. Weights fitted in training subsets were applied to the full cohort for the displayed curves, yielding resubstitution summaries; held-out estimates used test participants only. Classifier comparisons used repeated stratified five-fold cross-validation with 10 repeats. These designs are labeled separately in the Methods and figure legends.

### Supplementary Note 6. Directional miRNA–mRNA definitions

The 19 miRNAs and 16 mRNAs form 304 possible pairs; 119 meet the primary target-support rule. Within-feature z scores define four relative-abundance states: miRNA-high/mRNA-low, miRNA-low/mRNA-high, co-high and co-low. The terms suppression and de-repression are directional labels for these configurations and do not establish target repression, co-packaging or regulation in recipient cells. Each participant's state burden is the fraction of eligible pairs in that state.

### Supplementary Note 7. Dataset roles and provenance

Three published resources extend discovery. Magen et al.28 provides total-plasma miRNA measurements and clinical follow-up; Eshima et al.21 provides bulk cortical expression; and Pineda et al.22 resolves cortical expression by cell class and state. These resources contain participants independent of discovery and of one another. Total plasma informed miRNA-panel refinement and prognostic evaluation. The cortical datasets informed mRNA-subset selection and within-tissue interpretation; they do not test correspondence between an individual blood measurement and cortical state. Accessions, sample counts and analytical units are given in the main Methods.

Cortical inference used donor-level biological replication. Profiles or nuclei from the same donor were not treated as independent replicates. Complete contrast-level results and unavailable or non-estimable entries are retained in the Source Data and Supplementary Data.

### Supplementary Note 8. Chronology and independence of plasma analysis

The 19-miRNA list was fixed from the GLAST-positive discovery cohort before the total-plasma data were interrogated. Plasma survival effects subsequently entered the ranking that selected the final five-miRNA panel. The plasma cohort therefore provides independent participants for transfer and refinement, but not a fully external validation of the final panel; that step requires prospective evaluation with feature identities, assay and score fixed in advance.

### Supplementary Note 9. Measurement properties across compartments

Detection, RNA diversity, interparticipant variation and serial stability were evaluated on the expression scales specified below. These measurements describe different properties of the profiled RNA.

The candidate miRNAs had lower interparticipant log2CPM coefficients of variation than the plasma background: medians were 17.2% versus 54.2% in ALS and 15.6% versus 19.7% in controls. In GLAST-positive sEVs, the corresponding values were 21.9% versus 17.4% and 19.2% versus 15.5%. At the log2CPM >5 threshold, ALS samples contained 220 ± 82 detected miRNAs in the sEV fraction and 542 ± 129 in plasma; median Shannon diversity was 4.20 and 5.26, respectively (Extended Data Fig. 7g–i). Coefficients of variation were calculated on log2CPM across participants; Figure 1h reports a separate feature-richness analysis.

Serial plasma measurements in 22 participants provide feature-specific stability summaries for the five miRNAs and miR-181 comparators (Extended Data Fig. 7m). Stability varied across features. Supplementary Tables S3 and S4 report Cox-effect magnitudes and bootstrap direction concordance; S26 separately summarizes expression fold changes.

### Supplementary Note 10. Biomarker context and intended use

| Attribute | Statement |
|---|---|
| **Biomarker category** | Prognostic |
| **Context of use** | Candidate baseline prognostic tool for established ALS, evaluated alongside NfL and ALSFRS-R slope |
| **Population** | Participants with clinically manifest ALS |
| **Matrix** | Total plasma for proposed prognostic scoring; GLAST-positive sEVs for surface-selected discovery |
| **Measurand** | Composite abundance score for five plasma miRNAs: miR-31-5p, miR-4492, miR-93-3p, miR-421 and miR-324-5p; weighting is specified for each analysis. |
| **Analytical validation** | RNA-sequencing evaluation completed; targeted-assay development requires fixed qPCR/ddPCR specifications, precision estimates, detection and quantitation limits, and preanalytical and interlaboratory robustness. |
| **Clinical validation** | Retrospective transfer and panel refinement completed; prospective multisite evaluation requires a locked assay and score |
| **Scope** | Candidate baseline prognosis in established ALS; prospective evaluation is described in Note 10. |

The five-miRNA plasma panel is a candidate for baseline prognostic stratification in clinically manifest ALS, subject to prospective evaluation alongside NfL and ALSFRS-R trajectory. The paired mRNA candidates provide a route to biological investigation in separate cortical resources; they are not a validated blood measure of cortical state.

Prospective evaluation should measure calibration and incremental prediction against prespecified clinical and NfL baselines. The five feature identities, assay platform, normalization, coefficients, cut points and prediction horizons should be fixed before outcomes are analyzed. The prespecified separation of short- and longer-horizon performance is motivated by increasing evidence that circulating ALS biomarkers provide nonuniform information across disease time (main refs. 9, 15 and 16).

Samples should be randomized across processing batches and measured blind to outcome. Participant-level resampling should account for repeated observations or predictions. Trial applications could first evaluate the score as a prespecified covariate; treatment-response and enrollment uses would require evidence matched to those decisions.

### Supplementary Note 11. Cortical lineage, pathway and literature context

Supplementary Tables S8, S25 and S26 retain the post-nomination literature audit for the 35 candidates, the constraint and variant annotations used in the Fig. 4d sensitivity analysis, and

ALS-context evidence for the 25 lineage-gated cortical programs. Citation labels S1–S28 refer to the Supplementary references below.

The audit distinguishes direct main-text evidence from supplementary screening-list mentions and related-family reports. Direct glial or neurodegeneration context was identified for 18 of the 35 nominated features; absence from this scoped audit is recorded as no confirmed hit and not as evidence of no biological relationship.

The Fig. 4d genetics component primarily reflects human constraint and variant annotation. Candidate-level ALS GWAS and Open Targets fields were zero in the frozen record and are not presented as ALS genetic validation. Program-level citations in Supplementary Table S26 provide disease context for the cortical analyses while preserving the distinction between contextual precedent and evidence generated in this study.

### Supplementary tables and data files

Supplementary Tables S1–S26 are supplied as one indexed workbook, and panel-specific Source Data contain the observations underlying each figure. Citation labels S1–S28 used in Supplementary Tables S8, S25 and S26 correspond to the reference list below.
